# Importin α Characterizes a Micronuclear Environment Associated with Genomic Instability in Human Cancer Cells

**DOI:** 10.1101/2023.06.06.543979

**Authors:** Yoichi Miyamoto, Reo Kisanuki, Rieko Oshima, Shige H. Yoshimura, Mutsumi Yokota, Kazumitsu Maehara, Chiaki Hata, Taro Tachibana, Hiroshi Kimura, Masato Koike, Yasuyuki Ohkawa, Toyomasa Katagiri, Yoshihiro Yoneda, Masahiro Oka, Hisato Saitoh

## Abstract

Micronuclei (MN) are membrane-enclosed chromatin bodies and hallmarks of genome instability. Here, we report that importin α, a key nuclear transport factor, is highly concentrated in a distinct subset of MN in cultured human cancer cells. This selective localization is not governed by classical nuclear transport pathways. Live-cell photobleaching revealed remarkably reduced mobility of importin α between MN and cytoplasm. In addition, the subset of importin α-positive MN exhibited collapsed nuclear envelopes and compromised barrier functions. Importin α was also enriched in euchromatin regions, where it colocalized with chromatin-regulating molecules. Importantly, importin α and DNA repair/sensing molecules such as RAD51, RPA2, and cGAS showed mutually exclusive localization in MN, indicating that MN comprise distinct internal environments. These findings identify importin α as a molecular marker of the restricted MN state, representing a previously unrecognized microenvironment distinct from subsets characterized by conventional molecular markers of disrupted MN. This framework provides new insights into how MN heterogeneity underlies genome instability and immune evasion during cancer progression.

**Summary statement:** Accumulation of importin α in micronuclei, followed by modulation of the microenvironment of the micronuclei, suggests a non-canonical function of importin α in genomic instability and cancer development.

## INTRODUCTION

Micronuclei (MN) are cytoplasmic structures embedded within the nuclear envelopes but spatially separated from primary nuclei (PN). MN arise from lagging chromosomes or chromosome fragments produced by mitotic errors or DNA damage after genotoxic stress (Hintzsche et al, 2017; Kwon et al, 2020). Thus, MN are biomarkers of genome instability, including forms such as chromosome rearrangements, mutagenesis, and chromothripsis (Krupina et al, 2021; Kwon et al, 2020; Zhang et al, 2015). Recent studies further demonstrated that the structural fragility of MN and aberrant DNA replication within them provide mechanistic insights into the occurrence of catastrophic genome rearrangements, including chromothripsis (Crasta et al, 2012; Ly et al, 2017; Umbreit et al, 2020; von Appen et al, 2020). In particular, the study of chromothripsis in many aggressive types of cancer has been rapidly expanding and advancing. Research suggests that MN actively participate in chromothripsis during cancer development, progression, and genome evolution (Forment et al, 2012; Ly & Cleveland, 2017; Storchová & Kloosterman, 2016). Moreover, the exposure of DNA/chromatin fibers within MN to cytoplasmic components can serve as a potent molecular pattern signal by activating sterile inflammatory responses, which are associated with many chronic age-associated conditions (Miller et al, 2021). The nuclear envelope of MN can be partially or completely disrupted, allowing cytoplasmic DNA sensors, such as cyclic GMP–AMP synthase (cGAS), to access micronuclear DNA and trigger innate immune responses via the cGAS–STING pathway (Harding et al, 2017; Li & Chen, 2018; Mackenzie et al, 2017).

Signal-dependent protein transport occurs via nuclear pore complexes embedded in the nuclear envelope (Fernandez-Martinez & Rout, 2021; Hampoelz et al, 2019; Knockenhauer & Schwartz, 2016). Importin α is a classical nuclear localization signal (cNLS) receptor that acts as an adaptor molecule between cNLS-bearing cargo and its carrier molecule, importin β1 (Goldfarb et al, 2004). The cNLS–importin α–importin β1 trimeric complex migrates into the nucleus via mediation by the Ras-like small GTPase, Ran, and dissociates upon the binding of importin β1 and RanGTP (Christie et al, 2016; Wente & Rout, 2010). Importin α is exported to the cytoplasm by cellular apoptosis susceptibility gene product (CAS), also known as chromosome segregation 1-like protein [CSE1L; (Behrens et al, 2003; Stewart, 2007)]. Human express seven importin α subtypes, including importin α1 (KPNA2), importin α3 (KPNA4), importin α4 (KPNA3), importin α5 (KPNA1), importin α6 (KPNA5), importin α7 (KPNA6), and importin α8 (KPNA7), which exhibit distinct cNLS-binding specificities and selective expression in various tissues (Goldfarb et al, 2004; Miyamoto et al, 2012; Pumroy & Cingolani, 2015). Importin α1/KPNA2 has been well investigated, and its high expression levels are associated with poor prognosis in various cancers (Han & Wang, 2020; Sun et al, 2021; Wang et al, 2012). Notably, a strong nuclear staining of importin α1 in breast cancer cells using immunohistochemical analysis is substantially associated with a higher tumor stage (Alshareeda et al, 2015; Dahl et al, 2006; Dankof et al, 2007).

In addition to nucleocytoplasmic transport regulation, importin α plays non-transport roles in spindle assembly regulation, lamin polymerization, nuclear envelope formation, protein degradation, stress response, gene expression, cell surface response, and mRNA metabolism, which emphasizes its multi-functionality (Miyamoto et al, 2016; Oka & Yoneda, 2018). Importin α accumulates in the nucleus under exceptional conditions, such as stress responses, where it binds to a DNase-sensitive nuclear component, indicating its potential to associate with DNA/chromatin (Furuta et al, 2004; Kodiha et al, 2008; Miyamoto et al, 2004). We have previously demonstrated that nuclear-localized importin α associates with chromatin and regulates gene expression (Yasuda et al, 2012). A recent report suggested that the importin β binding (IBB) domain enables the binding of importin α to DNA (Jibiki et al, 2021). These findings indicate that nuclear-localized importin α can function as a regulator of nuclear homeostasis through its intrinsic capacity for DNA/chromatin binding.

In this study, we discovered that importin α was markedly concentrated in MN. Our findings indicate that the micronuclear localization of importin α does not fully conform to its conventional role and dynamics as a nuclear transport factor. Importantly, our results indicate that importin α delineates a subset of MN that differs from those defined by conventional markers of disrupted MN, pointing to an alternative microenvironmental state. Collectively, our findings suggest that importin α can serve as a molecular marker to reveal the structural and functional diversity of MN in human cancer cells.

## RESULTS

### Visualization of importin **α** accumulation in MN

Importin α is highly expressed in various cancer cells, and there is a potential association between its high expression levels and tumor stage and patient survival outcomes (Christiansen & Dyrskjot, 2013; Han & Wang, 2020). We validated the enhanced expression of importin α subtypes in breast cancer cell lines (MCF7, MDA-MB-231, and SK-BR-3) and a control, non-transformed human mammary cell line (MCF10A). Western blot assays revealed that importin α, particularly importin α1 (KPNA2), was expressed at higher levels in cancer cell lines than in the control (Fig. 1A). Next, we performed indirect immunofluorescence (IF) analysis on human cancer cell lines, including MCF7 and HeLa cells. Notably, we found that importin α1 accumulated prominently in MN located within the cytoplasm (MCF7 cells, Fig. 1B; HeLa cells, Fig. 1C; yellow arrowhead). The MN fluorescence intensity was considerably higher than that of primary nuclei (PN) or the cytoplasm, to the extent that the latter was almost invisible when normalized against the former (Fig. 1D). We also found that other importin α subtypes—importin α3 (KPNA4), importin α4 (KPNA3), and importin α5 (KPNA1)—were present in MN (Fig. 1E). Furthermore, similar MN subsets contained different subtypes, such as importin α3 or importin α5, along with importin α1 (Fig. S1A).

**Figure 1.**
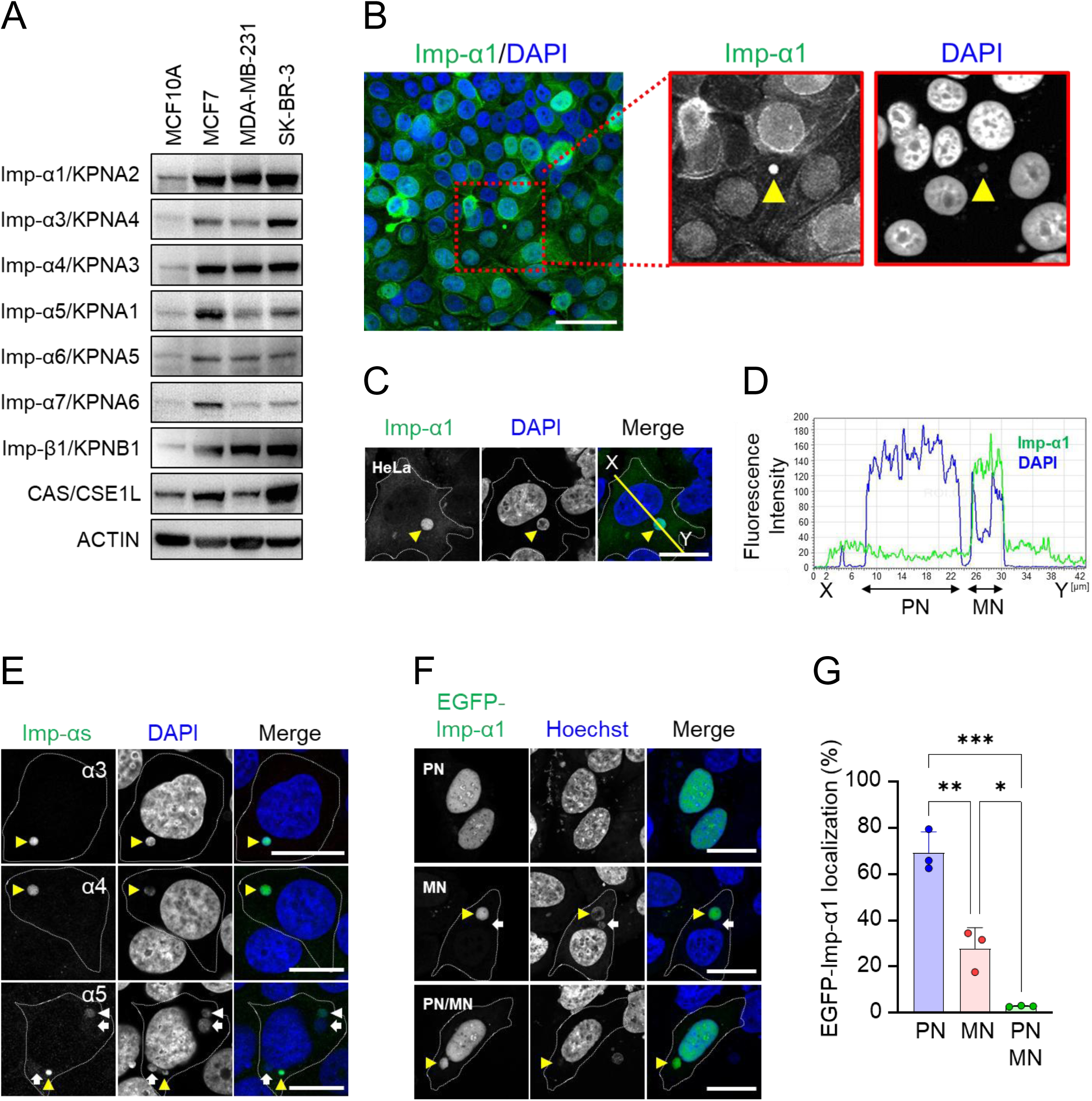
Accumulation of importin α proteins in MN. (A) Western blot of whole cell extracts from a human breast epithelial cell line (MCF10A) and breast cancer cell lines (MCF7, MDA-MB-231, and SK-BR-3) using antibodies against importin α (Imp-α/KPNA) subtypes, importin β1 (Imp-β1/KPNB1), CAS/CSE1L, and ACTIN, which served as a loading control. An antibody against importin α1 (KPNA2) was used as Ab2, as shown in Table S3. (B) Indirect IF images of MCF7 cells using an anti-importin α1 antibody (Imp-α1). The right panels show magnified views of the areas enclosed by the red dotted boxes in the left image. Yellow arrowhead indicates accumulation of importin α1 in the MN. DNA is stained with DAPI. Scale bar: 30 μm. (C) Indirect IF images of HeLa cells stained with anti-importin α1 antibody. Yellow arrowhead indicates MN with strong importin α1 signals. White dotted lines indicate cellular boundaries. DNA is stained with DAPI. The yellow line (X–Y) indicates the position of the fluorescence intensity profile shown in panel D. (D) Fluorescence intensity profiles of importin α1 (green) and DAPI (blue) plotted along the X–Y line shown in panel C. PN: primary nucleus, MN: micronucleus. (E) Indirect IF of MCF7 cells stained with anti-importin α3, α4, and α5 antibodies. Yellow arrowheads, white arrowheads, and white arrows indicate MN with high, low, or no importin α signal, respectively. Scale bar: 20 μm. (F) Subcellular localization of overexpressed EGFP–importin α1 (EGFP-Imp-α1) in HeLa cells treated with reversine for 20 h. PN: primary nuclei (upper panels); MN: micronuclei (middle panels); both PN and MN (PN/MN, lower panels). DNA is stained with Hoechest33342. Yellow arrowheads and white arrows indicate MN with either high intensity or no intensity of EGFP–Imp-α1 signals. Scale bar: 20 μm. (G) Percentage of EGFP–Imp-α1 localization in PN, MN, or PN/MN (corresponding to panel F). Data are shown as mean ± SD from three independent experiments (n = 3; approximately 35 cells per experiment; 108 cells in total). Statistical significance is determined using one-way ANOVA followed by Tukey’s multiple comparison test. *p < 0.05, **p < 0.01, ***p < 0.001.

Since MN are known to arise from lagging chromosomes during anaphase, a process associated with chromosome instability, which is a hallmark of cancer cells, we treated cells with an MPS1 inhibitor, reversine, which efficiently induces chromosome mis-segregation and MN formation [Fig. S2A, S2B (Agustinus et al, 2023; D’Alise et al, 2008; Santaguida et al, 2010)]. By treating HeLa cells with a range of reversine concentrations, we identified 400 nM for 20 h as the most effective condition for inducing MN formation without causing detectable cytotoxicity (Fig. S2C, S2D). To quantify the induction of MN by 400 nM reversine, we calculated the percentage of MN relative to the number of DAPI-stained PN in each microscopic field. In the untreated 542 cells analyzed across three independent experiments, MN formation rate was 5.5% ± 1.4. In contrast, reversine treatment significantly increased the MN formation rate to 42.0% ± 8.3. These results demonstrate that reversine robustly induced MN formation in HeLa cells without detectable cytotoxicity.

Indirect IF analysis using an anti-importin α1 antibody also revealed that a subset of MN exhibited importin α1 accumulation (Fig. S2E, positive: yellow arrowhead, negative: white arrow). We then examined whether reversine treatment affected the proportion of importin α1-positive MN. The results revealed that the MN formation rate for either untreated or treated cells was 36.2% ± 7.8 or 38.3% ± 8.8, respectively, with no significant difference (Fig. S2F). We also analyzed the frequency of MN formation and the proportion of importin α1-positive MN in non-transformed MCF10A cells and transformed MCF7 and MDA-MB-231 cells. Under untreated conditions, MCF7 and MDA-MB-231 cells exhibited a higher frequency of MN than MCF10A cells; however, this difference was no longer observed following reversine treatment (Fig. S3A). Furthermore, the proportion of importin α1-positive MN remained unchanged regardless of reversine treatment (Fig. S3B). These results imply that the spontaneous formation of MN is elevated in transformed cells, but importin α1 accumulation in MN is independent of transformation status.

We then examined MN localization of the overexpressed importin α1 fused with enhanced green fluorescence protein (EGFP) or monomeric red fluorescence protein (mRFP) in reversine treated HeLa cells. Notably, overexpressed importin α predominantly localizes in PN (Miyamoto et al, 2020). Consistent with this finding, over 60% of the EGFP–importin α1 expressing cells localized to the PN region (Fig. 1F, 1G). In contrast, a distinct pattern was observed in approximately 30% of the cells, where EGFP–importin α1, like its endogenous counterpart, was found in only a subset of MN. Localization to both MN and PN area simultaneously was very infrequent, detected in fewer than 3% of cells (Fig. 1F, 1G). Moreover, mRFP–importin α1 colocalized with EGFP–importin α subtypes (α3, α4, and α5) in the same MN (Fig. S1B), indicating that the accumulation of importin α family members in MN is a common feature.

### MN localization of importin **α**1 is uncharacterized in the context of conventional nuclear transport machinery

To determine whether the nuclear transport factors known to influence the subcellular localization of importin α coordinate its localization in importin α1-positive MN, we conducted an indirect IF analysis of importin β1, CAS/CSE1L, and Ran in reversine-treated HeLa cells. Importin β1 was predominantly observed in the cytoplasm and at the rim of PN, as well as in MN. However, most importin β1 signals represented the rim of the MN, with minimal overlap of importin α1 signals from inside the MN (Fig. 2A, top panels). Notably, approximately 17% of MN displayed importin β1 localization (Fig. 2B), proportion markedly lower than that of importin α1, and the co-localization rate with importin α1 was below 40% (Fig. 2C). These findings suggest that the selective and characteristic MN localization of importin α1 cannot be fully explained by its association with importin β1. In contrast, CAS/CSE1L and Ran signals were detected in the majority of MN, with localization rates of up to 88% or 91%, respectively (Fig. 2A, 2B; CAS/CSE1L: middle panels, Ran: bottom panels). Unlike importin β1, these factors were broadly distributed across MN, and their signals substantially overlapped with those of importin α1 (Fig. 2C). Taken together, these results indicate that CAS/CSE1L and Ran do not account for the selective MN localization of importin α1, and therefore, this localization does not conform to the conventional dynamics of a nuclear transport factor.

**Figure 2.**
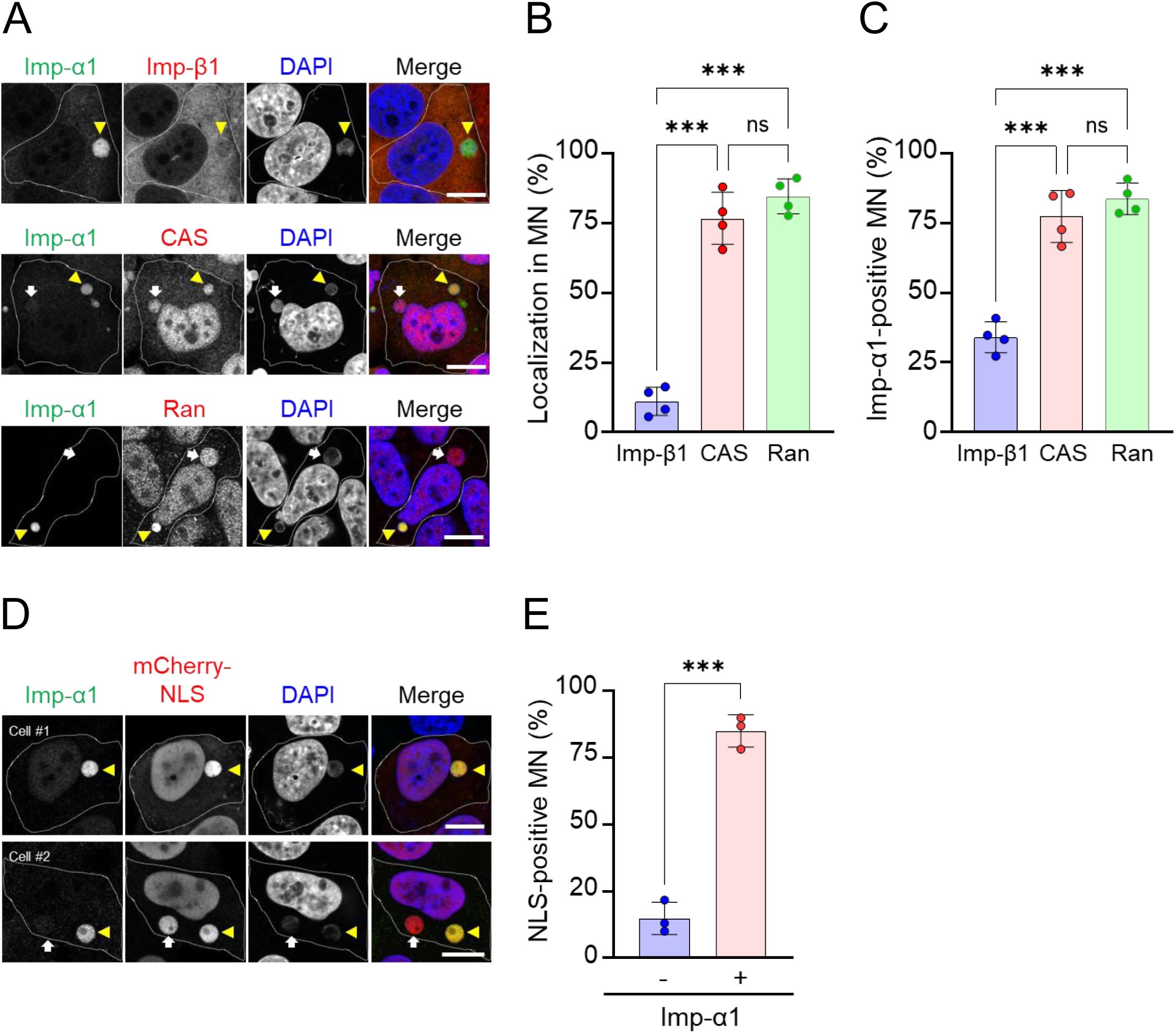
Localization of nuclear transport factors and cNLS substrates in importin α1-enriched MN. (A) Indirect IF images of importin α1 with importin β1, CAS/CSE1L, or Ran in reversine-treated HeLa cells. DNA is stained with DAPI. White dotted lines outline the cellular boundaries. Yellow arrowhead indicates high-intensity importin α1 signals in MN, whereas the white arrow indicates no importin α1 signals. Scale bar: 10 μm. (B) Percentage of MN positive for importin β1, CAS/CSE1L, or Ran in reversine-treated HeLa cells, as shown in panel A. Data are shown as mean ± SD from four independent experiments (n = 4; 80 MN in total for each condition). Statistical significance is determined using one-way ANOVA followed by Tukey’s multiple comparison test. ***p < 0.001; ns, not significant. (C) Percentage of MN colocalized with importin α1 and importin β1, CAS/CSE1L, or Ran in reversine-treated HeLa. Data are shown as mean ± SD of cells from four independent experiments (n = 4; 80 MN in total). Statistical significance is determined using one-way ANOVA followed by Tukey’s multiple comparison test. ***p < 0.001; ns, not significant. (D) Indirect IF images of mCherry–NLS-expressing HeLa cells treated with reversine, showing MN with or without importin-α1 localization. Yellow arrowheads indicate Imp-α1-positive MN, and white arrows indicate Imp-α1-negative MN. Scale bar: 10 μm. (E) Percentage of NLS-positive MN with or without importin α1 localization in reversine-treated HeLa cells expressing mCherry–NLS, as shown in D. Data are shown as mean ± SD from four independent experiments (n = 4, 170 MN in total). Statistical significance is determined using an unpaired two-tailed t-test. ***p < 0.001.

We then investigated whether MN localization of importin α1 was related to the transport of cNLS substrates. HeLa cells expressing mCherry–SV40TNLS contained endogenous importin α1 with a relatively high probability in cNLS-positive MN (Cell #1 yellow arrowhead in Fig. 2D, 2E). However, cNLS-signals were also detected in importin α1-negative MN (Cell#2, white arrow in Fig. 2D, 2E), indicating that the selective MN localization of importin α1 is not necessarily linked to the transport of cNLS substrates into MN.

To confirm that the MN localization of importin α1 does not occur due to the interaction with importin β1, CAS/CSE1L and the cNLS-substrates, we tested several importin α1 mutants, including those with the IBB domain deleted (ΔIBB), deficient in cNLS binding (ED), or mutated at the CAS/CSE1L binding domain (C-mut) to express an EGFP-fused protein (Fig. S4A). All the tested mutants were clearly observed in the MN (Fig. S4B, yellow arrowheads). In addition, different patterns of PN localization were identified. In contrast to wild-type importin α1, which appeared to be quite clear and localized exclusively in MN, C-mut localized in both MN and PN (Fig. S4B, S4C). To further clarify this observation, we quantified the MN/PN fluorescence ratio (Fig. S4D). To standardize nuclear size differences, fluorescence intensities were measured within a fixed circular area (1.5–2.0 µm in diameter), as illustrated by the red circles in the schematic above. This normalization allowed direct comparison of the signal levels between MN and PN. The ΔIBB mutant was also observed in PN, but to a lesser extent than C-mut. These findings suggest that the subcellular dynamics of importin α1 in PN and MN are quite different; MN localization of importin α1 is regulated by mechanisms distinct from those governing PN localization, which is typically associated with canonical nucleocytoplasmic recycling processes, including CAS/CSE1L-mediated shuttling of importin α1. Taken together, our data suggest that MN localization of importin α does not entirely conform to its traditional role as a nuclear transport factor.

### Remarkably low mobility of importin **α**1 in the MN

To further determine whether the localization of importin α1 in MN reflects regulation by canonical nuclear transport pathways, we assessed its dynamic properties by performing fluorescence recovery after photobleaching (FRAP) experiments in reversine-treated HeLa cells expressing EGFP–importin α1, in which both the PN and MN were fully photobleached within the same cells (Fig. 3A, 3C). These analyses revealed that the fluorescence recovery of importin α1 was significantly delayed in MN compared to PN, indicating a much slower nuclear import and/or recycling rate in MN.

**Figure 3.**
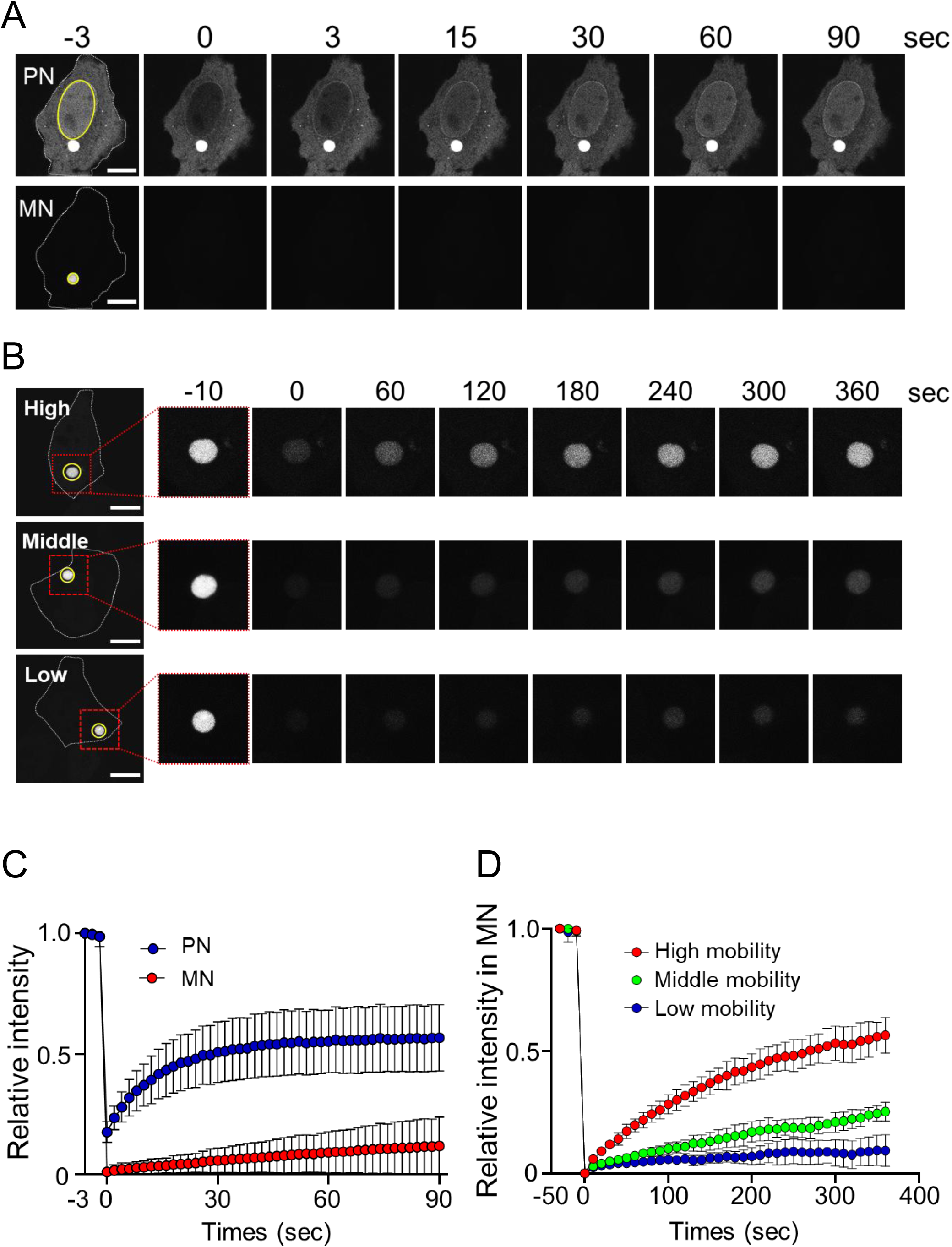
FRAP analysis of EGFP–importin α1 in MN. (A) Time-lapse images of EGFP–importin α1 in reversine-treated HeLa cells at the indicated time points before and after photobleaching. The image contrast has been individually optimized in PN and MN to clarify the signal loss and recovery. The entire principal nucleus (PN, top) and micronucleus (MN, bottom) are bleached within the same cell. The bleached area is indicated by a yellow circle. Scale bar: 10 μm. (B) Time-lapse images of EGFP–importin α1 in MN at the indicated time points. The bleached area encompassing the entire MN is indicated by a yellow circle. The right panels show magnified views of the region enclosed by the red dotted box in the left image. Three recovery patterns are shown: high, middle, and low. Scale bar: 10 μm. (C) Quantification of FRAP analysis in panel A. Fluorescence recovery of EGFP–importin α1 is plotted as normalized intensity (%) at the indicated time points after bleaching. (D) Quantification of FRAP analysis in panel B. Fluorescence recovery of EGFP–importin α1 in MN is plotted as normalized intensity (%) at the indicated time points. MN are classified into three groups based on the level of EGFP–importin α1 accumulation: high, middle, and low. Data are shown as mean ± SD from six MN (n = 6 per group).

Based on this observation, we further investigated the heterogeneity of importin α1 dynamics in MN by conducting detailed FRAP analyses across multiple MN (Fig. 3B, 3D). Importin α1 recovery profiles were classified into three distinct categories: high, intermediate (middle), and low mobility. These findings suggest that MN exhibit variable and generally reduced importin α1 mobility, possibly reflecting altered or impaired nuclear transport machinery within the MN. Taken together, these results demonstrate that importin α1 displays remarkably different trafficking dynamics in MN relative to PN, supporting the view that MN harbor distinct nuclear transport and recycling mechanisms

### Nuclear envelopes are partially collapsed in the importin **α1**-enriched MN

Because MN formation is frequently associated with nuclear envelope defects that can perturb nucleocytoplasmic trafficking (Guo et al, 2020; Hatch et al, 2013; Maciejowski & Hatch, 2020), we examined the nuclear envelope of the importin α1-enriched MN to further characterize their structural features using antibodies against lamins B1 and A/C. Indirect IF analysis revealed that neither lamin correlated with the selective MN localization of importin α1 (Cell #1 in Fig. 4A, 4B). Instead, importin α1 tended to concentrate in MN regions deficient in lamins (Cell #2 in Fig. 4A, 4B: see the graphs of the fluorescence intensity in the X–Y lines).

**Figure 4.**
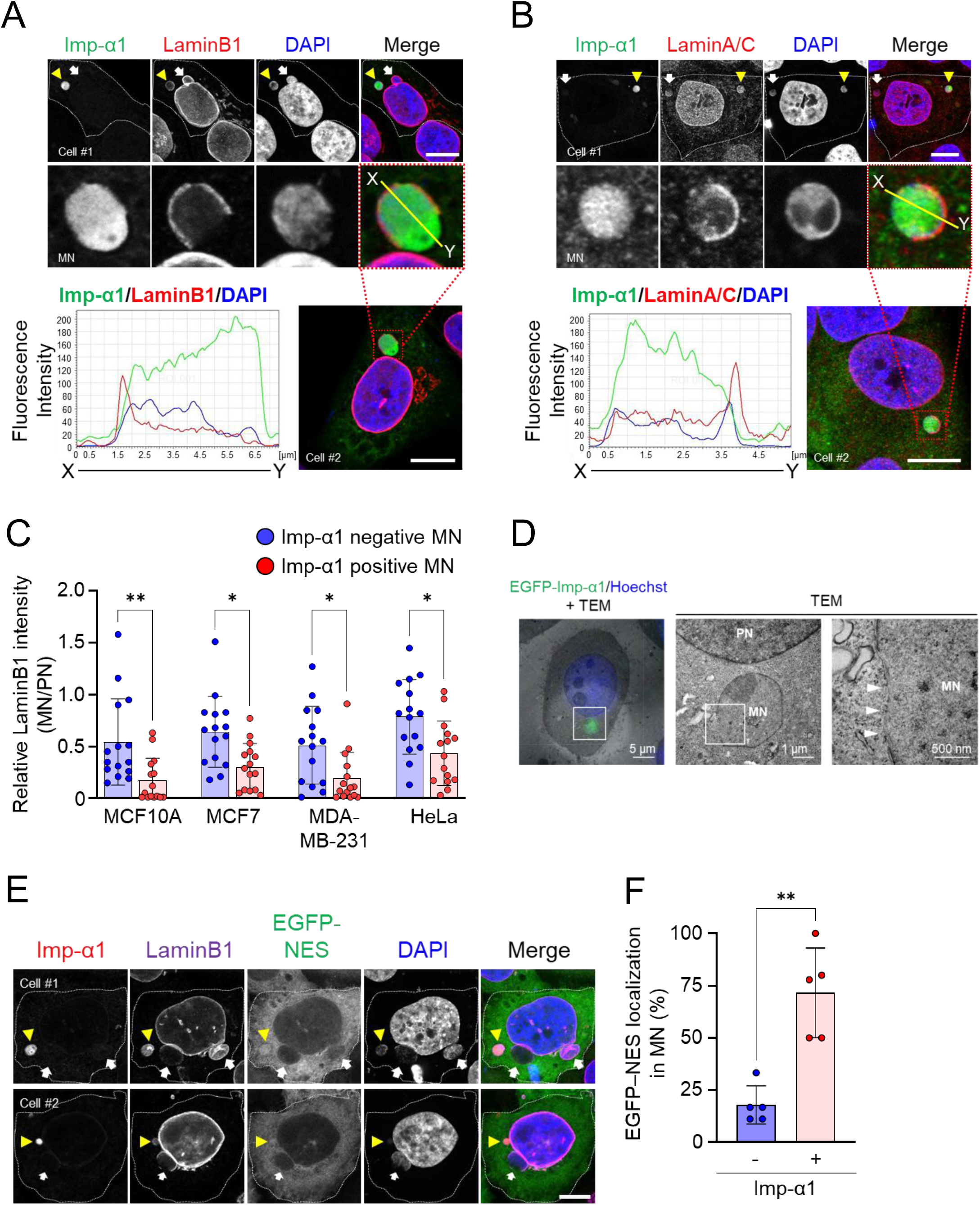
Nuclear envelope collapse of importin-α1-enriched MN. (A–B) Indirect IF images of importin α1 together with lamin B1 (A) or lamin A/C (B) in reversine-treated HeLa cells. Upper panels: Cell #1 (top, low magnification) and Cell #2 (bottom, high magnification with an X–Y line). Lower panels: the right image shows a low-magnification view of Cell #2, where the red dotted box corresponds to the enlarged image in the upper panels (bottom). The left image shows the fluorescence intensity profiles of importin α1 (green), lamin B1 or lamin A/C (red), and DAPI (blue) along the X–Y line in Cell #2. Yellow arrowhead indicates strong importin α1 signals, whereas white arrow indicates no importin α1 signals. Scale bar: 10 μm. (C) Relative LaminB1 intensity (MN/PN) in importin α1-positive and -negative MN in MCF10A, MCF7, MDA-MB-231, and HeLa cells, quantified from indirect IF images shown in panel A. For each cell, LaminB1 fluorescence intensity in MN is normalized to that of PN in the same cell. Each data point represents one cell (15 cells in total from three independent experiments). Data are shown as the mean ± SD. Statistical significance is determined using two-way ANOVA followed by Sidak’s multiple comparisons test. *p < 0.05, **p < 0.01. (D) CLEM analysis of importin α1-expressing MN in reversine-treated HeLa cells. The left panel shows merged EGFP–importin α1, Hoechst 33342, and TEM images; the boxed area indicates the EGFP-positive MN. Scale bar: 5 µm. The central panel shows an enlarged TEM image of the MN in the left box (box area). Scale bar: 1 µm. The right image shows an enlarged TEM image of the box region in the central panel. White arrowheads indicate areas lacking both inner and outer nuclear membranes in the MN. Scale bar: 500 nm. (E) MN localization of importin α1 and lamin B1 in EGFP–NES-expressing HeLa cells treated with reversine. Images of two independent cells are shown in the upper (Cell #1) and lower panels (Cell #2). Yellow arrowheads indicate Imp-α1-positive MN, whereas white arrows indicate Imp-α1-negative MN. Scale bar: 10 μm. (F) Percentage of MN with EGFP–NES in importin α1-negative and -positive MN. Data are quantified from approximately 10 cells per microscopy field, with five independent fields analyzed (approximately 35 cells in total from four independent experiments). Data are shown as mean ± SD (n = 5). Statistical significance is determined using a paired two-tailed t-test; **p < 0.01.

We assessed lamin B1 localization in importin α1-positive MN across four human cell lines (MCF10A, MCF7, MDA-MB-231, and HeLa) using indirect IF analysis. Lamin B1 signal intensity in MN was measured relative to that in the corresponding PN of the same cell, and the values were compared between importin α1-positive and -negative MN. This analysis revealed that, irrespective of whether the cells were non-transformed or cancer-derived, lamin B1 localization was consistently and significantly reduced in importin α1-positive MN compared to importin α1-negative MN in all tested cell lines (Fig. 4C). These quantitative results validate that nuclear envelope integrity is reduced in importin α1-positive MN, consistent with a tendency toward partial disruption of the micronuclear membrane.

To further validate this interpretation, we performed correlative light and electron microscopy (CLEM) on HeLa cells expressing EGFP–importin α1. Transmission electron microscopy (TEM) revealed portions of the nuclear envelope with marked weakening (indicated by white arrowheads in Fig. 4D).

We then employed an EGFP-fused nuclear export signal (NES) to assess the nuclear envelope in importin α1-positive MN. In HeLa cells expressing EGFP–NES, EGFP signals were excluded from the importin α1-negative MN (Fig. 4E, white arrows), whereas they originated from the importin α1-positive MN (Fig. 4E, yellow arrowheads). Statistical analysis revealed that the fluorescence intensity of the EGFP–NES signal was remarkably higher in importin α1-positive MN (Fig. 4F), indicating that the nuclear envelopes were partially collapsed. These results indicate that the nuclear envelopes were partially disrupted in a subset of importin α1-positive MN.

### Epigenetic chromatin status and importin **α** distribution in MN

In addition to nuclear envelope alterations, the chromatin environment may contribute to the unique features of importin α1-positive MN. Based on our previous findings that importin α can associate with DNase-sensitive chromatin (Yasuda et al, 2012), we assessed the relationship between importin α1 and DNA/chromatin status in MN by examining well-characterized histone H3 modifications. We used tri-methylations at lysine 4 and 9 of histone H3 (H3K4me3 and H3K9me3) as markers of euchromatin and heterochromatin regions, respectively. Indirect IF analysis showed that importin α1 signals positively correlated with H3K4me3 signals and negatively correlated with H3K9me3 signals in MDA-MB-231 cells (Fig. 5A, 5B), MCF10A cells (Fig. S5A, S5B), MCF7 cells (Fig. S5C, S5D) and HeLa cells (Fig. S5E, S5F). Consistent with these observations, we quantified the co-localization of importin α1 with H3K4me3 or H3K9me3 in the MN of MCF10A, MCF7, MDA-MB-231, and HeLa cells. Statistical analysis revealed that importin α1 was significantly higher in regions marked by H3K4me3 than in those marked by H3K9me3 across all cell lines (Fig. 5C), indicating that importin α1 preferentially localizes to euchromatic rather than heterochromatic regions within the MN.

**Figure 5.**
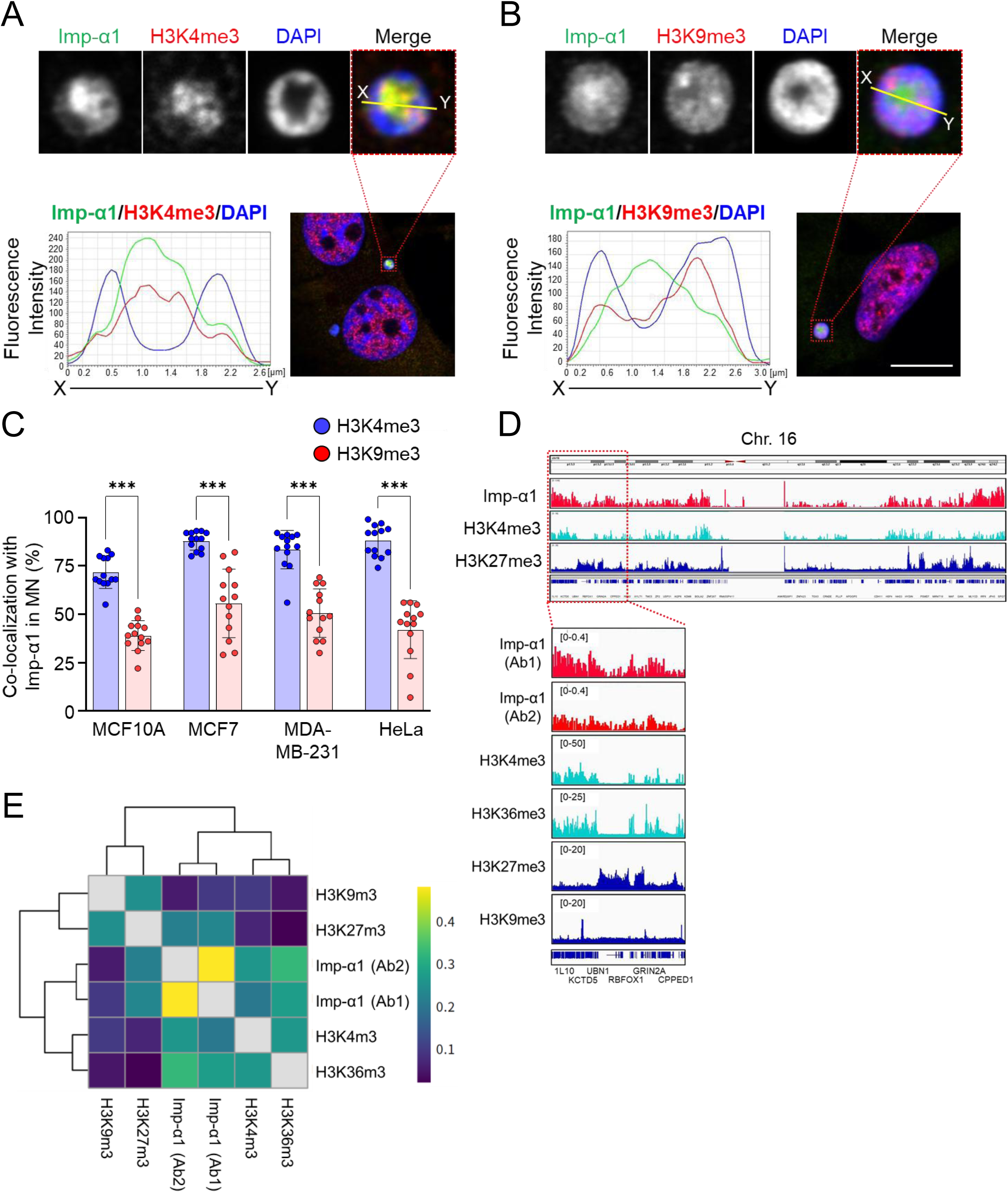
Euchromatin localization of importin α1 in MN. (A–B) Indirect IF images of importin α1 with H3K4me3 (A) or H3K9me3 (B) in MDA-MB-231 cells under normal culture conditions. The upper panels show magnified views of the areas indicated by the red dotted boxes in the lower image. DNA is stained with DAPI. Fluorescent intensities of importin α1 (green), modified histones (red: H3K4me3 or H3K9me3), and DAPI (blue) in MN are plotted along the X–Y line and shown on the left. Scale bar: 10 µm. (C) Percentage of MN colocalized with importin α1 and H3K4me3 or H3K9me3 in reversine-treated MCF10A, MCF7, MDA-MB-231, and HeLa cells, quantified from indirect IF images shown in panels A and B. Data are shown as mean ± SD from three independent experiments (n = 13 MN per cell line). Statistical significance is determined using two-way ANOVA followed by Sidak’s multiple comparisons test; ***p < 0.001 for H3K4me3 vs. H3K9me3 in all cell lines. (D) Alignment of ChIP-seq data for importin α1 and modified histones in MCF7 cells, shown for chromosome 16. ChIP was performed using two independent importin α1 antibodies (Ab1 and Ab2; Table S3), and modified histones data were obtained from public datasets of the ENCODE project (H3K4me3: ENCSR000DWJ, H3K27me3: ENCSR000EWP, H3K36me3: ENCSR000EWO, H3K9me3: ENCSR000EWQ). (E) Spearman’s correlation coefficient matrix from ChIP-seq data using two independent importin α1 antibodies and modified histones H3K4me3, H3K36me3, H3K27me3, and H3K9me3 in MCF7 cells.

To explore the chromatin-binding activity of importin α1, we conducted chromatin immunoprecipitation sequencing (ChIP-seq) analysis on whole MCF7 cells using two different importin α1 antibodies. Comparison of the ChIP-seq data for importin α1 with a commercially available dataset for modified histones revealed that importin α1 is associated with euchromatin regions characterized by H3K4me3 and H3K36me3, but not with heterochromatin regions marked by H3K27me3 and H3K9me3 (Fig. 5D: represented chromosome 16, and Fig. 5E).

Taken together, our data indicate a preferential distribution of importin α1 in the euchromatic areas rather than in the heterochromatic regions of the MN.

### Search for potential importin **α**1-interacting proteins accumulated in MN

Given that importin α1 is concentrated in euchromatin regions, likely due to its chromatin-binding capacity, we attempted to identify potential importin α1-interacting partners. To achieve this, rapid immunoprecipitation mass spectrometry of endogenous proteins [RIME (Mohammed et al, 2016)] was performed on whole-cell extracts from untreated wild-type MCF7 cells using an anti-importin α1 antibody. Using two independent samples from MCF7 cells, we identified 102 proteins as potential importin α1 interactors, excluding 4 overlapping proteins (106 proteins, as shown in Fig. 6A and Table S1). Gene ontology analysis using the PANTHER Pathways database (Mi et al, 2013; Thomas et al, 2022) revealed that many of these proteins are involved in DNA replication and p53 pathways related to the DNA damage response (Fig. 6B).

**Figure 6.**
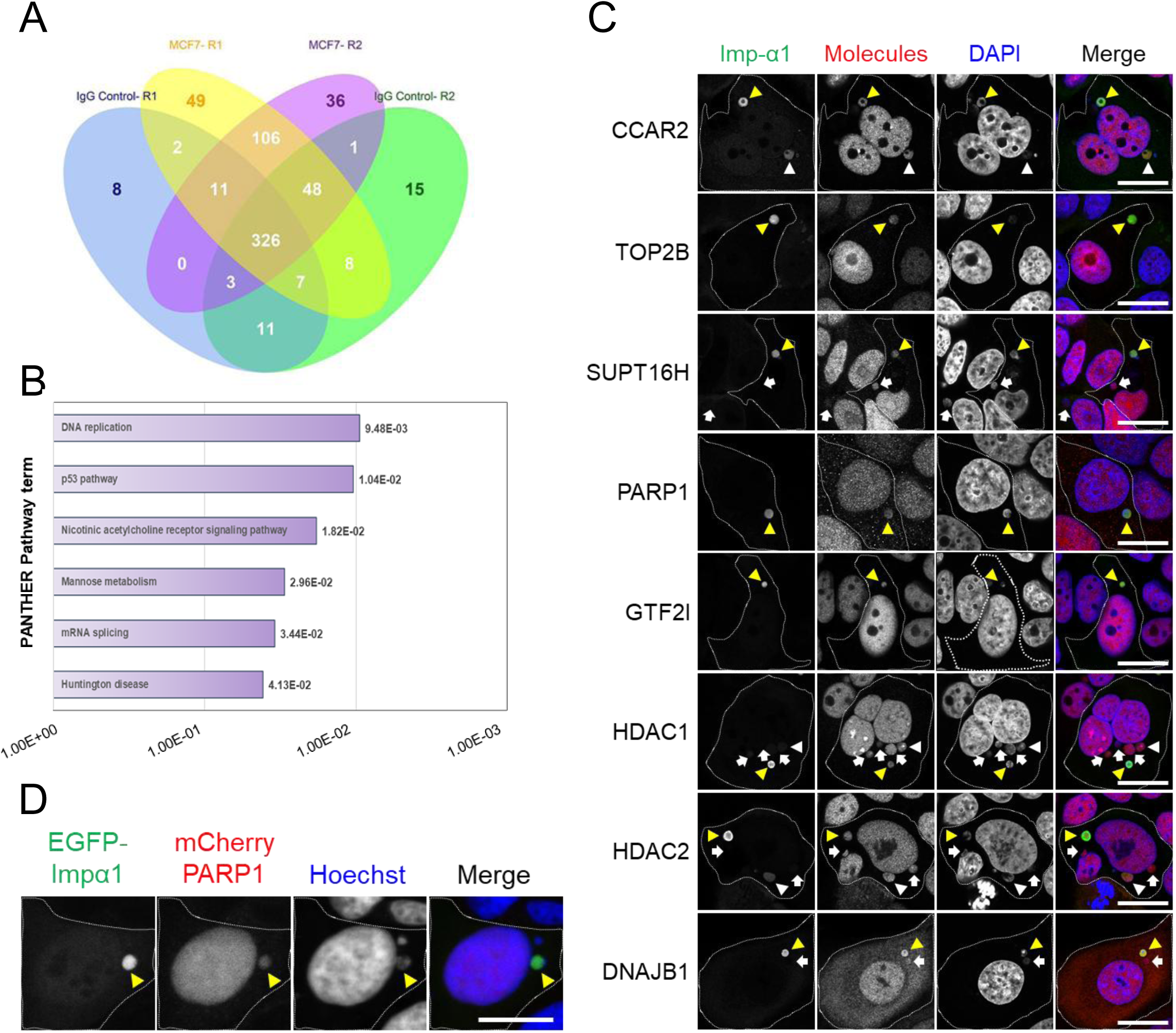
Exploration of importin α1-binding proteins using RIME. (A) Venn diagram of candidate importin α1-binding proteins identified using RIME. Data were obtained from two independent experiments. MCF7-R1 and MCF7-R2 represent proteins precipitated with anti-importin α1 antibodies, whereas IgG Control-R1 and -R2 represent proteins precipitated with normal IgG antibodies. A total of 106 proteins are identified as candidate binding partners of importin α1. (B) Gene Ontology (GO) analysis of the binding proteins identified by RIME. GO terms were obtained from the PANTHER Pathways database and ranked according to their p-values. (C) Indirect IF images of importin α1 with candidate molecules identified by RIME in MCF7 cells treated with 400 nM reversine for 20 h. Candidate molecules are detected using specific antibodies (Table S3). White dotted lines indicate cellular boundaries. DNA is stained with DAPI. Merged images are shown on the right. Yellow arrowheads indicate MN with high importin α1 signals, white arrowheads indicate MN with low signals, and white arrows indicate MN with no signals. Scale bar: 20 μm. (D) Co-localization of overexpressed EGFP–importin α1 with mCherry–PARP1 in MCF7 cells. Yellow arrowheads indicate MN containing both proteins. Scale bar: 10 µm.

As importin α1 interactors are typically nuclear proteins, it is plausible that they reside not only in the primary nucleus but also in the MN. To test this possibility, we screened the identified candidates for MN localization using immunofluorescence images provided by the Human Protein Atlas (HPA) database (Pontén et al, 2008; Thul et al, 2017). A careful review of the IF images in the HPA dataset revealed 60 proteins that appeared in both MN and PN. Of the 20 candidate MCF7 cell proteins identified by IF analysis, eight proteins—CCAR2/DBC1, TOP2B, SUPT16H, PARP1, GTF2I, HDAC1, HDAC2, and DNAJB1—were detected within MN (Fig. 6C). Notably, the signals for CCAR2/DBC1, TOP2B, SUPT16H, and PARP1 significantly overlapped with importin α1 signals from the MN (Table S2). Additionally, the mCherry–PARP1 fusion protein colocalized with EGFP–importin α1 in MN (Fig. 6D). These results highlight the chromatin-associated proteins that colocalize with importin α in MN.

### Mutually exclusive MN localization of importin **α**1 and RAD51

In our investigation of importin α1-intereacting partners and their roles in MN, we discovered that RAD51, a homologous recombination repair protein involved in the DNA damage response, accumulated in the MN of etoposide-treated HeLa cells (Fig. 7A). Up to 43% of MN exhibited RAD51-positive signals. Although most RAD51-positive MN displayed low-intensity signals (Fig. 7B, upper panels: Cell #1, white arrowhead), a small subset of MN occasionally showed significantly stronger fluorescence intensity (Fig. 7B, lower panels: Cell #2, yellow arrowhead).

**Figure 7.**
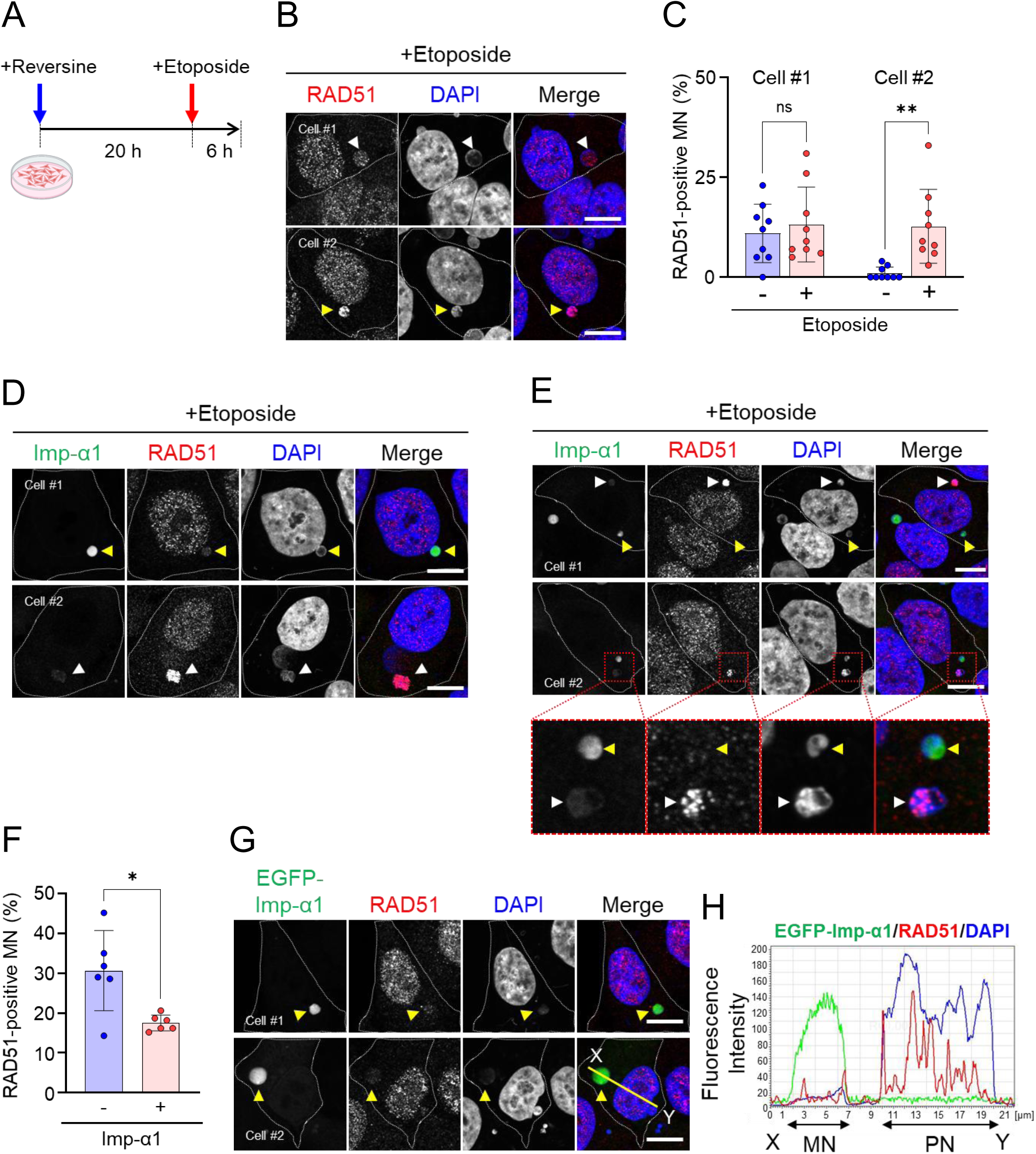
Mutually exclusive localization of importin α1 and RAD51 in MN. (A) Schematic representation of HeLa cell treatment with reversine, followed by etoposide. Cells are treated with 400 nM reversine for 20 h and then incubated with 10 µM etoposide for 6 h. (B) Indirect IF images of RAD51 in HeLa cells treated with 400 nM reversine for 20 h, followed by 10 µM etoposide for 6 h. Two independent cells are shown: the upper panels (Cell #1) contain MN with low RAD51 signal intensity (white arrowhead), and the lower panels (Cell #2) contain MN with high RAD51 signal intensity (yellow arrowhead). Scale bar: 10 μm. (C) Percentage of RAD51-positive MN in HeLa cells treated with reversine with or without etoposide. MN are classified according to RAD51 signal intensity as low (Cell #1) or high (Cell #2), as exemplified in panel B. Data are from nine fields (n = 9; 302 MN untreated, 213 MN etoposide-treated), with each dot representing one field. Data are shown as the mean ± SD. Statistical significance is determined using two-way ANOVA followed by Sidak’s multiple comparisons test (**p < 0.01; ns, not significant). (D) Indirect IF images of importin α1 (green) and RAD51 (red) in HeLa cells treated with reversine followed by etoposide. Two independent cells are shown: the upper panels (Cell #1) contain MN with high importin α1 and low RAD51 signals (yellow arrowhead), whereas the lower panels (Cell #2) contain MN with low importin α1 and high RAD51 signals (white arrowhead). Scale bar: 10 μm. (E) Indirect IF images of importin α1 (green) and RAD51 (red) in HeLa cells treated with reversine followed by etoposide. The upper panels (Cell #1) show a cell outlined by a white dotted line containing two MN: one with high importin α1 signal (yellow arrowhead) and the other with high RAD51 signal (white arrowhead). The middle panels (Cell #2) show two MN in a single cell. The lower panels present enlarged views of the areas enclosed by the red dotted boxes. Yellow arrowheads indicate MN with high importin α1 and low RAD51 signals, whereas white arrowheads indicate MN with low importin α1 and high RAD51 signals. Scale bar: 10 μm. (F) Percentage of RAD51-positive MN, comparing importin α1-negative and -positive MN in HeLa cells, as exemplified in panels D and E. A total of 191 importin α1-negative MN and 190 importin α1-positive MN were analyzed across six microscopy fields (n = 6) in three independent experiments. Data are shown as the mean ± SD. Statistical significance is determined using a paired two-tailed t-test (*p < 0.05). (G) Indirect IF images of RAD51 in HeLa cells overexpressing EGFP–importin α1 (EGFP–Imp-α1). Cells are transfected with EGFP–Imp-α1 and subsequently treated with reversine and etoposide. Two independent cells are shown (Cell #1 and #2). DNA is stained with DAPI. Yellow arrowheads indicate EGFP–Imp-α1 localized MN. The yellow X–Y line in the lower panels (Cell #2) indicates the position used for the fluorescence intensity profile shown in panel H. Scale bar: 10 μm. (H) Fluorescence intensities of EGFP–Imp-α1 (green), RAD51 (red), and DAPI (blue) in MN plotted along the X–Y line indicated in panel G.

Based on this difference in RAD51 signal intensity, we classified RAD51-positive MN into Cell #1 (low-intensity) and Cell #2 (high-intensity) and quantified their frequencies under untreated and etoposide-treated conditions (Fig. 7C). The analysis revealed that etoposide treatment markedly increased the number of Cell #2-type MN, indicating that RAD51 accumulates strongly in a specific subset of MN following DNA damage induction. These findings suggest that etoposide preferentially enhances RAD51 accessibility in high-intensity-type MN.

Notably, we found that the localization of importin α1 and RAD51 in the MN was mutually exclusive. MN with low RAD51 signal intensity exhibited a strong importin α1 signal intensity (Fig. 7D, upper panels: Cell #1, yellow arrowhead), and vice versa (Fig. 7D, lower panels: Cell #2, white arrowhead). Moreover, these molecules exhibited distinct compartmentalization while localizing to MN in a single cell (Fig. 7E, Cells #1 and #2: yellow arrowhead; importin α1-positive and RAD51-negative MN, white arrowhead; importin α1-negative and RAD51-positive MN). Quantitative analysis showed that RAD51-highly positive MN (Cell#2 type in Fig. 7B) were significantly less frequent among importin α1-positive MN than among importin α1-negative MN (Fig. 7F). These results support the mutually exclusive relationship between importin α1 accumulation and RAD51 localization in the MN.

Finally, we found that the signal intensity of endogenous RAD51 was remarkably low in MN overexpressing EGFP–importin α1 (Fig. 7G, 7H). Importantly, such an altered distribution was not observed for one of the binding candidates, PAPR1(Fig. 6D), indicating that the mutual exclusion was specific to RAD51.

### Importin **α**1-positive MN lack DNA recognition molecules RPA2 and cGAS

Following our finding that RAD51 and importin α1 localize in a mutually exclusive manner within MN, we further analyzed replication protein A2 (RPA2/RPA32), a core subunit of the RPA complex that coats ssDNA and facilitates the subsequent recruitment and filament assembly of RAD51, thereby ensuring homologous recombination repair (Bhat & Cortez, 2018; Krejci et al, 2012). Based on this functional connection, we investigated whether RPA2 localizes to MN in relation to importin α1. We found that, even more prominently than RAD51, RPA2 was largely absent from importin α1-positive MN and instead localized to a distinct subset of MN that lacked importin α1 accumulation (Fig. 8A).

**Figure 8.**
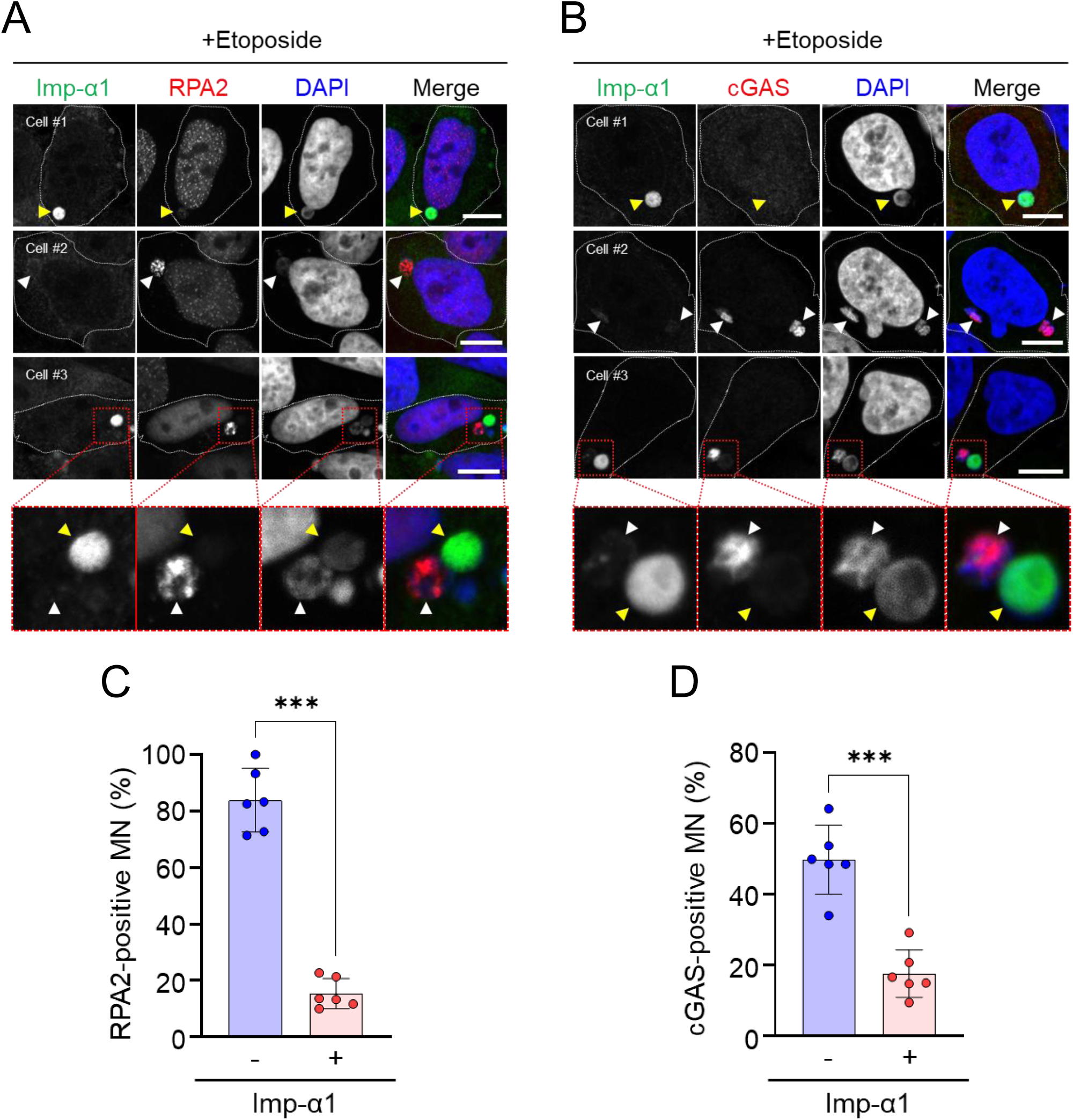
Importin α1 localization is mutually exclusive to RPA2 and cGAS in MN. (A–B) Indirect IF images of HeLa cells treated with reversine followed by etoposide, stained for importin α1 (green) together with RPA2 (A, red) or cGAS (B, red), and DAPI (blue). Three representative cells are shown: Cell #1 (top) contains an importin α1-positive MN lacking RPA2 or cGAS signals; Cell #2 (middle) contains an importin α1-negative MN with RPA2 or cGAS localization; Cell #3 (bottom) contains two MN in a single cell, one importin α1-positive MN and the other RPA2- or cGAS-positive MN. The lower panels show magnified views of the areas enclosed by the red dotted boxes. Yellow arrowheads indicate importin α1-positive MN, and white arrowheads indicate RPA2- or cGAS-positive MN. Scale bars: 10 μm. (C–D) Percentage of RPA2-positive MN (C) and cGAS-positive MN (D) among importin α1-negative and importin α1-positive MN in HeLa cells, as exemplified in panels A and B. For RPA2, a total of 191 importin α1-negative MN and 135 importin α1-positive MN are analyzed across six fields (n = 6). For cGAS, 230 importin α1-negative MN and 140 importin α1-positive MN are analyzed across six fields (n = 6). Data are shown as the mean ± SD. Statistical significance is determined using paired two-tailed t-tests (***p < 0.001).

Given that both RAD51 and RPA2 were excluded from importin α1-positive MN, we investigated whether this selective pattern also applies to cytosolic DNA sensors such as cyclic GMP–AMP synthase (cGAS). cGAS is a well-characterized sensor that gains access to micronuclear DNA upon rupture and is linked to chromothripsis, genome instability, and innate immune signaling (Harding et al, 2017; Ly et al, 2017; Mackenzie et al, 2017; Vietri et al, 2020; von Appen et al, 2020). Therefore, we analyzed the localization of cGAS in relation to importin α1. cGAS showed a distribution similar to that of RPA2, being rarely detected in importin α1-positive MN and instead accumulating in importin α1-negative MN (Fig. 8B). Statistical analyses further supported that importin α1-positive MN and importin α1-negative MN constitute distinct subsets that are differentially accessible to DNA sensor molecules (RPA2: Fig. 8C; cGAS: Fig. 8D).

Taken together, these results indicate that DNA-recognizing molecules such as RAD51, RPA2, and cGAS preferentially localize to MN lacking importin α1 accumulation, whereas importin α1 defines a separate subset of MN. Thus, importin α1 and DNA-recognizing proteins exhibit distinct and largely mutually exclusive localization patterns depending on the MN microenvironment, highlighting importin α1 as a molecular marker that reflects a unique MN state.

## DISCUSSION

In this study, we found that importin α was localized in the MN of human cancer cells, indicating that MN localization is a common feature of the importin α family of proteins. Notably, importin α was localized in some MN but not in others. This observation suggests that the specific features of the MN state, such as its structure and function, may be related to the unique function of importin α.

MN localization of importin α1 did not fully correlate with the distribution of classical nuclear transport factors, such as importin β1, CAS/CSE1L, and Ran, or with the transport of cNLS substrates into the MN. All EGFP–importin α1 mutants (ΔIBB, ED, and C-mut), which lack canonical transport functions, showed MN localization. The C-mut, defective in nuclear export, was detected in both PN and MN, whereas wild-type importin α1 was observed exclusively in MN. Indeed, our quantitative analysis showed that the reduced MN/PN ratio in the C-mut resulted primarily from increased accumulation of importin α1 in the PN, rather than reduced retention in the MN. This finding supports that CAS/CSE1L-mediated export of importin α1 is active in the PN but may be impaired or functionally uncoupled in the MN. This functional discrepancy was further supported by FRAP experiments, in which both PN and MN were fully photobleached within the same cells. In particular, importin α1 is rapidly recycled from the PN, whereas it moves very slowly between the MN and cytoplasm. Together, these results provide strong evidence that the nucleocytoplasmic recycling of importin α1 is selectively collapsed in MN.

We demonstrated that importin α1 was enriched in lamin-deficient areas of MN, as evidenced by electron microscopy analysis, which showed a fragile nuclear envelope morphology. Therefore, we propose that importin α1 mediates the construction and/or collapse of the nuclear envelope of MN. Indeed, a previous study demonstrated that exogenously added recombinant importin α proteins inhibited nuclear envelope assembly around decondensed sperm chromatin in *Xenopus* egg extracts (Hachet et al, 2004), indicating that highly concentrated importin α inhibits nuclear membrane formation. An alternative scenario that cannot be ruled out is that importin α accumulation is not the cause of nuclear membrane collapse but rather a consequence of it. Recent studies have indicated that interactions between mitochondria and MN can induce nuclear envelope damage and/or inhibit the repair of nuclear membrane defects, possibly via reactive oxygen species generated by mitochondria located near MN (Di Bona et al, 2024; Martin et al, 2024). Although we did not collect data on the proximity of importin α-positive MN to mitochondria in this study, our findings suggest alternative mechanisms that may promote nuclear membrane damage or impede the repair of membrane defects in MN.

The preferential distribution of importin α1 in the euchromatin regions of MN provides a plausible explanation for its markedly reduced mobility. Our ChIP-seq analysis confirmed that importin α1 is enriched in chromatin regions marked by H3K4me3 and H3K36me3, further supporting its affinity for transcriptionally active domains. Consistent with this, previous studies have demonstrated that MN often undergo defective and asynchronous DNA replication, leading to replication stress, DNA damage, and chromosomal fragmentation (Crasta et al, 2012). Consistent with these findings, our RIME analysis identified replication stress–responsive factors, such as PARP1 and the FACT subunit SUPT16H, which were enriched in importin α1–positive MN and colocalized with importin α1 in a subset of cells. Importantly, a previous study demonstrated that nuclear-localized importin α associates with DNase-sensitive chromatin components and participates in gene regulation (Yasuda et al, 2012), reinforcing the notion that importin α1 interacts with chromatin-associated regulators. These data suggest that importin α1 is retained in MN through stable interactions with euchromatin-associated regulators that respond to replication stress. Such interactions likely contribute to the impaired nucleocytoplasmic recycling observed in MN and may define a chromatin-linked state of importin α1 that reflects the replication-associated stress environment of MN.

Our study uncovered a previously unrecognized relationship between importin α1 and DNA-recognizing molecules within MN. RAD51 showed mutually exclusive localization with importin α1, likely restricting its access to DNA in importin α1-positive MN. RPA2, an ssDNA-binding partner of RAD51, was also largely absent from importin α1-positive MN and instead accumulated in importin α1-negative MN. The exclusion of RAD51 and RPA2 from importin α1-positive MN implies a deficiency in homologous recombination repair, providing a mechanism by which these MN contribute to chromothripsis and genome instability. In addition, cGAS showed a similar distribution, being rarely observed in importin α1-positive MN but accumulating in importin α1-negative MN. Because cGAS–STING activation is linked to chromothripsis, genome instability, and innate immune signaling (Harding et al, 2017; Mackenzie et al, 2017; Vietri et al, 2020), its exclusion suggests that importin α1-positive MN are inaccessible to immune surveillance. Importantly, our findings indicate that importin α1 and cGAS are targeted to MN via distinct mechanisms. Importin α1 preferentially localizes to H3K4me3-enriched regions, consistent with a chromatin-associated mode of binding, whereas cGAS binds primarily to non-chromatinized DNA exposed by nuclear envelope rupture (Hatch et al, 2013; Sun et al, 2013). Notably, in our indirect IF analysis, MN positive for cGAS, RAD51 and RPA2, often displayed irregular morphologies, further supporting the idea that these molecules recognize non-chromatinized DNA in disrupted MN. Together, these results indicate that MN can be broadly categorized into at least two characteristic environments: one in which genomic DNA is exposed and accessible to repair and sensing molecules such as RAD51, RPA2, and cGAS, and another in which strong importin α accumulation correlates with their exclusion and relative genomic preservation. These categories likely represent predominant tendencies rather than strictly binary states, and additional intermediate or context-dependent variations may exist in the data. The distinction between these two categories further suggests that importin α defines a previously unrecognized subset of MN, distinct from the conventional ‘ruptured MN’ classically characterized by cGAS recruitment. The conceptual framework provides a new understanding of MN heterogeneity and highlights how the internal state of MN critically influences genomic stability, cell fate decisions, and ultimately cancer progression.

Notably, importin α is closely linked to tumorigenesis and poor prognosis (Christiansen & Dyrskjot, 2013; Han & Wang, 2020). Previous studies have largely attributed this association to its nuclear transport activity, with elevated importin α thought to promote the nuclear entry of oncogenic proteins, thereby conferring a growth advantage to tumor cells (Huang et al, 2013; Li & Chen, 2018; Zhou et al, 2021). In this study, we demonstrate that importin α accumulates in a distinct subset of MN, independent of its transport activity, thereby defining a micronuclear environment that contributes to genome instability and immune evasion. From this perspective, our findings may provide a new explanation for clinical observations from large cohort studies in breast, ovarian, and lung cancers, where high importin α expression has been linked to advanced stage, recurrence, and poor survival (Alshareeda et al, 2015; Dahl et al, 2006; Noetzel et al, 2012; Wang et al, 2012).

## MATERIALS AND METHODS

### Cell lines and culture conditions

Human cervical cancer cell line HeLa cells (RIKEN, Tsukuba, Japan), Human breast cancer cell line MCF7 (NIBN, JCRB 0134, Osaka, Japan), MDA-MB-231, SK-BR-3, and human breast epithelial cell line MCF10A (gifted from Satoshi Fujii, National Cancer Center, Japan) were maintained in following culture conditions: HeLa and SK-BR-3 cells, Dulbecco’s Modified Eagle’s Medium (DMEM; Sigma-Aldrich, St. Luis, MO, USA); MCF7 cells, Minimum Essential Medium Alpha (MEMα; Life Technologies, Carlsbad, CA) supplemented with 0.1 mM MEM Non-essential amino acids (NEAA; Life Technologies), 1 mM Sodium pyruvate (Life Technologies), and 10 µg/mL Insulin (Wako, Osaka, Japan); MDA-MB-231 cells, RPMI1640 (Sigma-Aldrich) with 2 mM L-Glutamine (Life Technologies), and MCF10A cells, MEG mammary epithelial cell growth medium (MEGM; added to SingleQuote (CC-3150). All media contained 10% fetal bovine serum (FBS). Cells were cultured at 37 °C in a 5% CO2 incubator. Reversine (Cayman Chemical, MI, USA) and etoposide (Tokyo Chemical Industry (TCI) Co., LTD., Tokyo, Japan) were used as described in the figures.

### Antibodies

The primary antibodies used are listed in Table S3. The secondary antibodies used were: Alexa Fluor Plus 488 (A32723, A32790, A32731, and A32814; Thermo Fisher Scientific Inc., Rockford, IL, USA) and Alexa Fluor 488 (A11055), Alexa Fluor 594 (A-21203, A32754, and A11032) for indirect IF and horseradish peroxidase (HRP)-conjugated anti-goat, anti-rabbit, or anti-mouse secondary antibodies (Jackson ImmunoResearch Inc West Grove, PA, USA) for western blotting.

### Indirect immunofluorescence analysis

Cells were cultured on 18 × 18 mm coverslips (Menzel-Glaser, Braunschweig, Germany) in a 35 mm dish and fixed with 3.7% formaldehyde in phosphate-buffered saline (PBS) for 15 min, followed by treatment with 0.5% Triton X-100 in PBS for 5 min. After blocking with 3% skim milk in PBS for 30 min at 25 °C, the cells on the coverslips were labeled with primary antibodies overnight and then reacted with secondary antibodies for 1 h at 25 °C. To visualize DNA, the slides were incubated with DAPI (1:5000 in PBS; Dojindo Laboratories, Kumamoto, Japan) for 20 min. Fluorescence images were obtained using a confocal microscope (Leica TCS SP8 II; Leica Microsystems, Wetzlar, Germany). A detailed description of the image analysis is provided in the Supplementary Materials and Methods.

### Plasmid construction for mammalian expression

Human importin α1 (KPNA2) cDNA subcloned into the pGEX6P2 plasmid was gifted by Akira Tsujii (NIBIOHN) and then cloned into the pEGFPC1 vector (Clontech, Mountain View, CA, USA). Plasmids for pEGFP-mouse importin α1 (KPNA2) were constructed as described previously (Yasuda et al, 2012). The SV40T antigen NLS (NLS; PKKKRKVED) within pmCherry-C1 (pmCherry–NLS) was constructed as previously described (Miyamoto et al, 2022). The pEGFPC1-NES plasmid was kindly provided by Dr. Asally (Asally et al, 2011). cDNAs encoding PARP1 and RAD51 were amplified from HeLa cells using PCR. Primer information is provided in the Supplementary Materials. cDNAs were cloned into the pmCherryC1 vector using the In-Fusion HD cloning kit (Clontech, Mountain View, CA, USA).

### Transfection

HeLa cells were plated onto 18 × 18 mm coverslips (Menzel–Glaser) in a 35 mm dish for 2 days prior to transfection. Transfection of mammalian expression plasmids was performed using Lipofectamine 2000 DNA Transfection Reagent (Thermo Fisher Scientific) following the manufacturer’s instructions.

### Western blotting

Western blotting was performed as previously described (Miyamoto et al, 2022). Whole cell lysates (20 μg) were subjected to 10% SuperSep SDS-PAGE gel (Wako, Osaka, Japan). The membranes were incubated with primary antibodies (dilutions ranging from 1:2000) diluted in Can Get Signal Immunoreaction Enhancer Solution 1 (TOYOBO, Osaka, Japan) overnight at 4 °C. The HRP-conjugated secondary antibodies (dilutions ranging from 1:8000 to 1:10,000) were diluted in Can Get Signal Immunoreaction Enhancer Solution 2 (TOYOBO) at 25 °C for 1 h. The signals were detected using an ATTO EzWest lumi plus (WSE-7120S, ATTO Corporation, Tokyo, Japan).

### Correlative light-electron microscopy (CLEM)

CLEM analysis was performed as described previously (Yokota et al, 2021), with some modifications. Briefly, HeLa cells attached on gridded coverslips (Matsunami, Osaka, Japan) were transfected with pEGFP-human importin α1 (KPNA2) for 1 day and treated with 400 nM reversine. The cells were then fixed in 2% paraformaldehyde, 0.5% glutaraldehyde, and 50 mM sucrose in 0.1 M phosphate buffer and stained with Hoechst 33342 for 1 h. After bright-field and fluorescence images were captured using a BZ-X810 microscope (KEYENCE, Osaka, Japan), the cells were further fixed in 2% glutaraldehyde and 50 mM sucrose in 0.1 M phosphate buffer and post-fixed with 1% osmium tetroxide. Fixed cells were dehydrated and embedded in Epok 812 (Oken Shoji, Tokyo, Japan). Ultrathin sections were cut using a UC6 ultramicrotome (Leica), stained with uranyl acetate and lead citrate, and examined using a JEM-1400 transmission electron microscope (JEOL).

### Chromatin immunoprecipitation sequence (ChIP-seq)

MCF7 cells (1 × 10^6^) were grown in 100 mm dishes (IWAKI, Tokyo, Japan) for 7 days and fixed at 25 °C for 5 min by adding 135 µL of 37% formaldehyde solution (Nacalai Tesque, Kyoto, Japan) in 10 mL of the culture medium. Fixed cells were washed twice with ice-cold PBS and collected with 0.5 mL of ice-cold PBS using a cell lifter (Corning, NY, USA) and centrifuged at 600 × *g* at 4 °C for 5 min. The cells were resuspended in ChIP buffer [10 mM Tris-HCl (pH 8.0), 200 mM KCl, 1 mM CaCl_2_, 0.5% NP40] containing a protease inhibitor cocktail (No.25955, Nacalai Tesque), briefly sonicated (Branson 250 Sonifier, Branson Ultrasonics), and incubated on ice for 10 min. After centrifugation at 10,000 × *g* at 4 °C for 10 min, the supernatants were digested with 1 U/mL micrococcal nuclease (Worthington Biochemical) at 37 °C for 30 min. The reaction was stopped with EDTA (final concentration of 10 mM) and centrifuged at 10,000 × *g* at 4 °C for 10 min. The supernatants were incubated with anti-rabbit or anti-mouse IgG magnetic beads (Dynabeads, Life Technologies) with anti-importin α1 (BD Biosciences, Cat. #610486; Abcam, Cat. # ab84440, Table S3) for 8 h. The beads were washed twice with ChIP buffer, ChIP wash buffer [10 mM Tris-HCl (pH 8.0), 500 mM KCl, 1 mM CaCl_2_, 0.5% NP40, a protease inhibitor cocktail], and TE buffer [10 mM Tris-HCl (pH 8.0) and 1 mM EDTA] and eluted overnight in an elution buffer containing [50 mM Tris-HCl (pH 8.0), 10 mM EDTA, and 1% sodium dodecyl sulfate (SDS)] at 65 °C. DNA was recovered using AMPure XP beads (Beckman Coulter) and used for ChIP-seq analysis.

A ChIP library was prepared using the NEBNext Ultra DNA Library Prep Kit for Illumina (New England Biolabs) following the manufacturer’s instructions, and sequencing was performed on an Illumina HiSeq1500 system. Sequence reads were aligned to the reference human genome (GRCh38) using Hisat2 (version 2.0.5). Genomic tracks (bigWig) and correlation matrix were generated using deepTools (version 3.4.5).

### Rapid immunoprecipitation mass spectrometry of endogenous proteins (RIME)

MCF-7 cells (1.5 × 10^6^) were cultured in 100 mm dishes (IWAKI) in 10 mL of MEM medium with 10% FBS for 5 days. To fix the cells, 1/10 volume (1 mL) of freshly prepared Formaldehyde Solution [FS: 11% Methanol-free formaldehyde, 0.1 M NaCl, 1 mM EDTA, and 50 mM Hepes (pH7.9)] was directly added to the medium and incubated for 8 min at 25 °C. After the addition of 0.55 mL of 2.5 M glycine solution to 11 mL of the medium (final conc. 0.1 M) to quench crosslinking for 5 min, and the cells were washed twice with ice-cold PBS. After removal of the supernatants, cells were collected at 800 × *g* for 10 min at 4 °C, and the cell pellets were resuspended in 10 mL of ice-cold PBS with 0.5% Igepal CA-630 (#I-8896, Sigma-Aldrich) and centrifuged again. The cell pellets were suspended in 10 mL of ice-cold PBS with 0.5% Igepal CA-630 and 100 µL of 100 mM phenylmethylsulfonyl fluoride (PMSF) in EtOH (final conc. 1 mM). The cell suspension was centrifuged thrice to pellet the cells, and the cell pellets were snap-frozen in liquid N2 and stored at −80 °C. RIME analysis was performed by Active Motif, CA., according to published protocols (Mohammed et al, 2016). RIME was performed using an antibody against importin α1 (KPNA2; Ab2 in Table S3) and 150 μg of chromatin from MCF7 cells to identify proteins that interact with importin α1 using mass spectrometry. Mass spectrometry of the immunoprecipitated protein complexes resulted in 57–60% coverage of importin α1.

### Fluorescence recovery after photobleaching (FRAP)

HeLa cells (ATCC, CCL-2.2) were cultured in DMEM supplemented with 10% FBS at 37 °C under 5% CO_2_. A plasmid encoding EGFP–human importin α1(KPNA2) was introduced into HeLa cells using polyethyleneimine (Polyscience, 49553937). The cells were replated onto a glass-bottom dish (D11130H, Matsunami) 12 h after transfection and incubated for 12 h. The cells were then incubated with reversine (400 nM) for 20 h. Fluorescence observation and FRAP analyses were performed using a confocal laser scanning microscope (Evident, FV3000). Time-lapse images of green fluorescence were obtained every 10 s for 6 min (MN) or every 3 s for 1.5 min (PN or MN). The fluorescence signal of the region of interest (ROI) was bleached by 488 nm laser irradiation. The fluorescence signals of the ROI were quantified using ImageJ (ver.1.54f).

### Accession number

ChIP-Seq data are accessible through the GEO Series with accession number GSE283000.

### Statistical analysis

Cells and MN were counted using a fluorescence microscope. Unless otherwise stated, the data are presented as the mean ± standard deviation (SD). The number of independent biological replicates (n) and total sample size are indicated in each figure legend. Statistical analyses were performed using GraphPad Prism version 9.0 (GraphPad Software, La Jolla, CA, USA). Comparisons between two independent groups were evaluated using unpaired two-tailed Student’s t-tests, whereas paired two-tailed t-tests were applied when comparing importin α1-positive and -negative MN within the same sample. For comparisons among more than two groups, one-way ANOVA with Tukey’s multiple comparison test was performed. For two-factor analyses (e.g., cell line × importin α1 status), two-way ANOVA with Sidak’s multiple comparison test was performed. Statistical significance was defined as p < 0.05, and significance levels are indicated in the figures as follows: p < 0.05 (*), p < 0.01 (**), p < 0.001 (***).

### Additional details

Extended experimental procedures (Animal care and use, antibody generation, plasmid construction, cell viability assays, detailed immunofluorescence analysis, and Quantification of MN) are described in the Supplementary Materials and Methods.

## ACKNOWLEDGMENTS

We thank Akira Tsujii (NIBIOHN) for kindly gifting the pGEX6P2-human importin α1 plasmid, Munehiro Asally (University of Warwick) for kindly gifting the pEGFPC1-NES plasmid, and Satoshi Fujii (National Cancer Center, Japan) for kindly gifting the human breast epithelial cells. We also thank the members of the Oka, Katagiri, and Saitoh Laboratories for their helpful discussions and technical support and the Laboratory of Morphology and Image Analysis, Biomedical Research Core Facilities, Juntendo University Graduate School of Medicine for technical assistance.

## AUTHOR CONTRIBUTIONS

YM and HS conceived the project and designed the experiments. YM, RK, RO, SHY, MY, KM, CH, TT, HK, MK, and YO conducted the experiments. YM, RK, SHY, MY, KM, MK, YO, TK, YY, MO, and HS analyzed the data and designed the figures. YM and HS drafted the manuscript.

