## Supplemental Data 1 for "Importin α Characterizes a Micronuclear Environment Associated with Genomic Instability in Human Cancer Cells"

#### *Supplemental Material*

### MATERIALS AND METHODS

##### Animal care and use

The experimental procedures for the production of monoclonal antibodies against importin α6 (KPNA5) were approved by the CEC Animal Care and Use Committee and performed according to the CEC Animal Experimentation Regulations.

##### Antibodies

A new mouse monoclonal antibody that specifically recognized importin α6 (KPNA5) protein was generated and named Clone number 1E3-2D6. As an antigen for immunization, bacterially produced recombinant importin α6 was purified as described previously (Miyamoto & Oka, 2016). An 8-week-old female C57BL/6 mouse (SLC, Shizuoka, Japan) was injected into the tail base with 100 µL of emulsion containing recombinant importin α6 protein and Freund's complete adjuvant. Briefly, 17 days after the first immunization, an additional immunization of the importin α6 protein was performed without an adjuvant into the tail base of the mouse. Four days after the additional immunization, cells from the iliac lymph nodes of the immunized mouse were fused with mouse myeloma Sp2/0-Ag14 cells at a ratio of 5:1 in 50% polyethylene glycol solution. The resulting hybridoma cells were plated onto 96-well plates and cultured in HAT selection medium. The importin α6-specific antibody was screened by ELISA, western blotting, and immunostaining using the hybridoma supernatants. Finally, a hybridoma clone producing a monoclonal antibody named 1E3-2D6 was selected. The Mab 1E3-2D6 was found to be an IgG 2a (k) subtype using a mouse isotyping kit.

The antibodies against histone modifications used for immunofluorescence analysis were anti-H3K4me3 (Kimura et al, 2008) and anti-H3K9me3 (Hayashi-Takanaka et al, 2011).

##### Plasmid construction and purification of recombinant protein

Mouse cDNAs of importin α1 (KPNA2), importin α3 (KPNA3), importin α4 (KPNA4), or importin α5 (KPNA1) in the pGEM-T vector (Tsuji et al, 1997) were subcloned into the pmRFPC1 or pEGFPC1 vectors (Clontech, Mountain View, CA, USA). Plasmids for pEGFP-mouse importin α1 (KPNA2) wild type and its mutants including the IBB domain-deleted mutant (termed ΔIBB mutant), the NLS-binding deficient mutant (D192K/E396R; termed ED mutant), and the CAS-binding defective mutant ((G469E/L470A/D471A/K472A/I473A/E474G; termed C-mut), were constructed as described previously (Yasuda et al, 2012).

The cDNAs encoding PARP1 and RAD51 were amplified from HeLa cells by PCR using the following primers: PARP1 forward: 5'-GTCCGGACTCAGATCTATGGCGGAGTCTTCGGAT-3' and 5'-CCGCGGTACCGTCGACTTACCACAGGGAGGTCTTA-3', and RAD51 forward: 5'-GTCCGGACTCAGATCTATGGCAATGCAGATGCAGCT-3' and reverse: 5'-CCGCGGTACCGTCGACTCAGTCTTTGGCATCTCCCACT-3'.

Purification of bacterially expressed FLAG-h-importin α1 (KPNA2) recombinant protein was performed as previously described (Miyamoto & Oka, 2016).

##### Cell viability assay

HeLa cells were plated in 96-well plates (SARSTEDT, Numbrecht, Germany) at 0.5–1 × $10^4$ cells per well for 24 h and then incubated with reversine for 20 h. Reversine was added at final concentrations of 0, 0.2, 0.4, 0.6, 0.8, 1, 2, 4, 6, 8, 10 µM. Cell viability was measured by Cell Counting Kit-8, (Dojindo Laboratories, Kumamoto, Japan) at the absorbance at 450 nm. n = 6

##### Immunofluorescence analysis

Fluorescence images were acquired using a Leica TCS SP8 confocal microscope equipped with an HC PL APO CS2 63×/1.40 NA oil immersion objective (Leica Microsystems, Wetzlar, Germany). Cells were sequentially excited with 405 nm, 488 nm, and 552 nm laser lines at 2–5% output power (measured at the back focal plane). The emission was collected in three channels using HyD-based spectral detectors and appropriate emission filters. The pinhole was set to 1 Airy Unit for each channel. Images were acquired as single optical sections without Z-stacking using the LAS X software (Leica Microsystems). The scan resolution was 1024×1024 pixels, with a zoom factor of 2.0–8.0×, yielding a pixel size of approximately 40–90 nm depending on the zoom. The scan speed ranged from 400 to 600 Hz. The detector gain and offset were optimized per channel to balance the signal-to-noise ratio (Heddleston et al, 2021).

All representative images shown in Figs 4A–B, 5A–B, and S5 are single optical sections. Similar discontinuities in the signals for lamin proteins (including laminB1 and laminA/C) and NPC components were consistently observed in multiple cells, indicating that these features were not attributable to image acquisition artifacts or to the membrane being located in another z-plane.

To quantify the frequency of micronuclei (MN; Supplemental Fig. S2D, S3A) and the proportion of importin α-positive MN (Supplemental Fig. S2F, S3B), and IF images were analyzed using ImageJ software (ver. 1.54f, NIH). Images stained with DAPI and anti-importin α antibodies were used for analysis. DAPI images were binarized by applying an appropriate intensity threshold, and overlapping nuclei were separated using the watershed algorithm. Particle analysis was performed using the "Analyze Particles" function with a size range of 100–infinity (pixels$^2$) and circularity set to 0.5–1.0 to eliminate debris and irregular structures. MN were defined as DAPI-positive particles smaller than primary nuclei, circular in shape, and located within the same cytoplasmic region as the associated nucleus. Cells with multiple nuclei or abnormal morphologies were excluded from the analysis. The proportion of cells containing at least one micronucleus was calculated for each field. For importin α staining, MN were categorized as importin α-positive if the fluorescence signal intensity clearly exceeded the surrounding cytoplasmic background signal. The percentage of importin α-positive MN was calculated among total MN per image. Quantification was performed across at least three biologically independent experiments, with 8–10 images analyzed per condition per replicate. Data are presented as the mean ± SD. Statistical comparisons were made using an unpaired two-tailed Student's t-test, one-way or two-way ANOVA (GraphPad Prism 9.0), with $p < 0.05$ considered statistically significant.

LaminB1 fluorescence intensity in MN and the corresponding primary nucleus (PN) within the same cell was quantified using Fiji (ImageJ, NIH; Fig. 4C). LaminB1 images were converted to binary masks using the Threshold function. The PN and MN within each cell were manually selected using the Oval Tool and registered as ROIs in the ROI Manager. For each ROI, the mean fluorescence intensity was measured from the original laminB1 image using the "Measure" function. The relative laminB1 intensity in each MN was calculated as the ratio of MN intensity to that of the PN in the same cell. Values were obtained from 15 cells per condition, collected across three independent experiments for statistical analysis.

To evaluate the degree of co-localization of importin α1 and histone modifications in MN (Fig. 5C), IF images were analyzed using Fiji (ImageJ, NIH). For each MN, a region of interest (ROI) was manually defined using the oval selection tool and registered in the ROI Manager. Binary masks for importin α1 and histone signals were generated via thresholding, and the overlapping region was calculated using the Image Calculator function. The area of each signal and their intersection was measured using the "Analyze Particles" function (size: 10–infinity). A total of 13 MN were analyzed per condition from three biologically independent experiments. Data are presented as mean ± SD. Statistical significance was assessed using Sidak's multiple comparisons test.

A

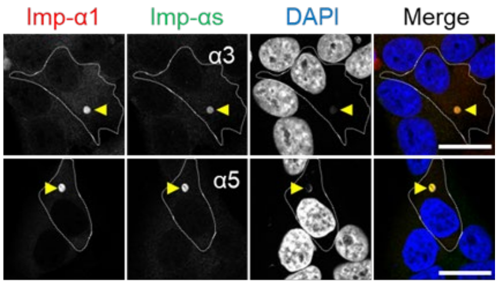


B

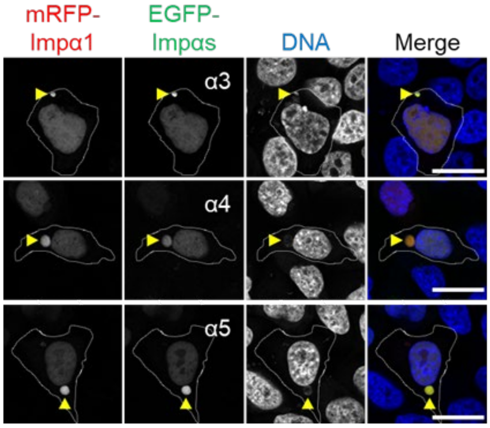


**Figure S1. Importin α subtypes localize to the same MN.** (A) Indirect IF images showing co-localization of importin α subtypes in MN of MCF7 cells. Importin α1 (red) co-localizes in the same MN with either importin α3 or importin α5 (green).Yellow arrowheads indicate MN with importin α subtype localization. Scale bar: 20 µm. (B) Co-localization of overexpressed mRFP–importin α1 with EGFP–importin α subtypes (α3, α4, and α5) in the same MN. Yellow arrowheads indicate MN with EGFP–importin αs. Scale bar: 20 µm.

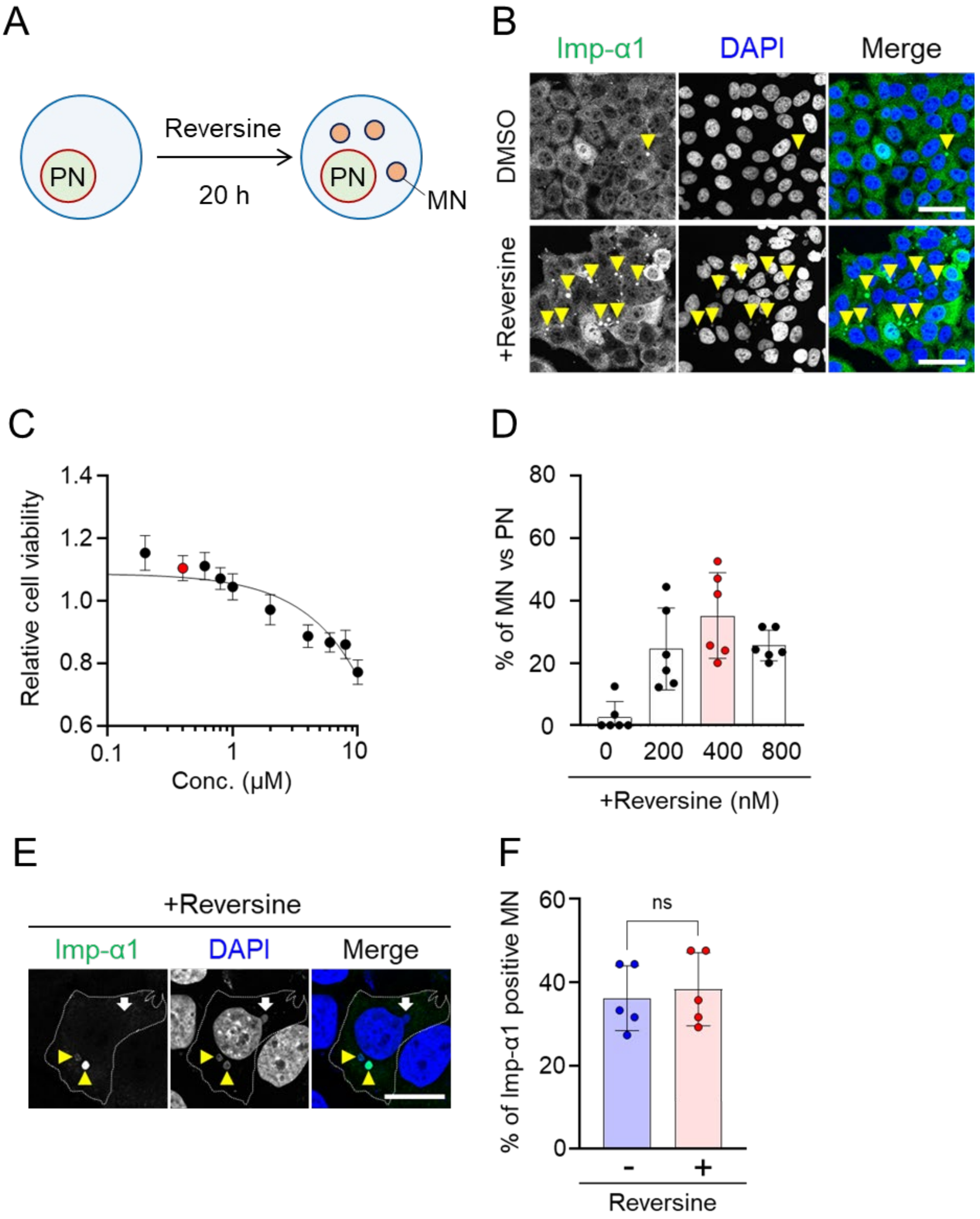


**Figure S2. MN localization of importin α in reversine-treated HeLa cells.** (A) Schematic representation of MN induction by reversine treatment of cultured HeLa cells. Cells are treated with reversine for 20 h. PN: primary nucleus; MN: micronuclei. (B) Indirect IF images of HeLa cells stained with anti-importin α1 antibody after treatment with DMSO (control) or 400 nM reversine for 20 h. Yellow arrowheads indicate MN positive for importin α1. Scale bar: 50 μm. (C) Relative cell viability of HeLa cells treated with the indicated concentrations of reversine. Cell viability is determined using the WST-8 assay. Data are shown as mean ±SD of six replicates from a single experiment, representative of three independent experiments. The 400 nM concentration is highlighted as a red point. (D) Ratio of MN number to PN number in HeLa cells under treated with reversine. Data are presented as mean ± SD from three independent experiments, each including six microscopy fields (n = 6). The 400 nM concentration is highlighted with a red bar. (E) Indirect IF images of HeLa cells treated with 400 nM reversine for 20 h and stained with anti-importin α1 antibody. Yellow arrowheads indicate importin α1-positive MN, and the white arrow indicates an importin α1-negative MN. Scale bar: 10 µm. (F) Quantification of importin α1 localization in MN of HeLa cells with or without reversine treatment. Data are shown as mean ± SD from three independent experiments, each including five microscopy fields (n = 5; untreated: 98 MN in total, reversine-treated: 99 MN in total). Statistical significance was determined using a paired two-tailed t-test.

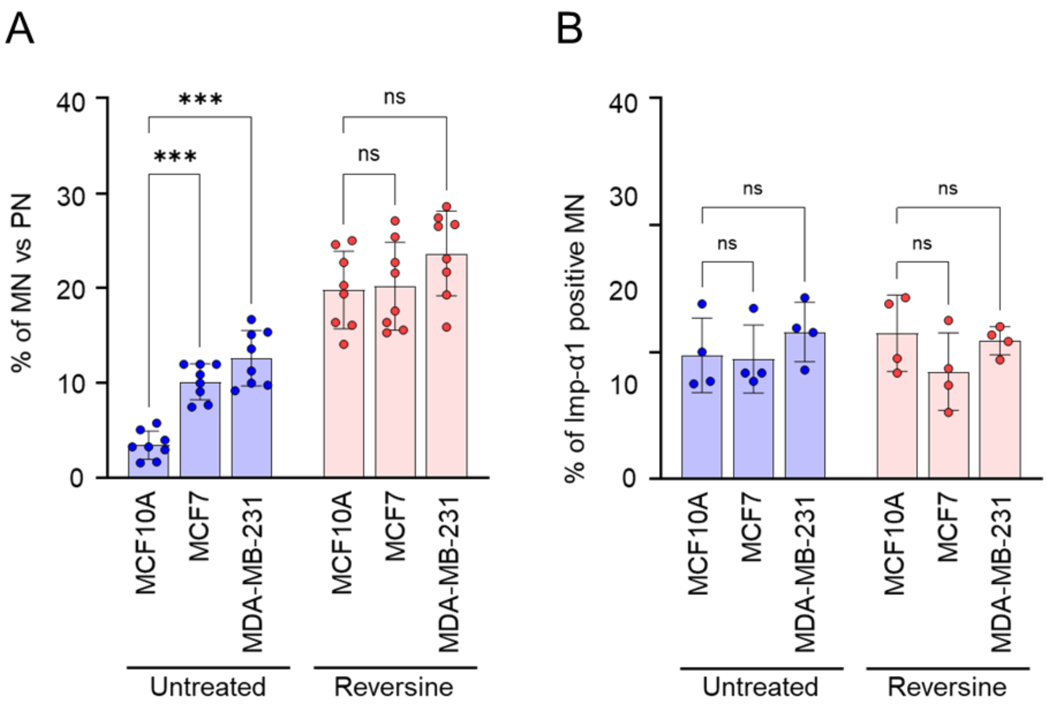


**Figure S3. Quantification of MN and importin α1-positive MN in human cell lines.**
(A) Ratio of MN to PN in MCF10A, MCF7, and MDA-MB-231 cells with or without reversine treatment. PN and MN are identified based on DAPI staining. Data are shown as mean ± SD from three independent experiments, each including eight microscopy fields (untreated: MCF10A, 471 cells in total; MCF7, 519 cells; MDA-MB-231, 465 cells; reversine: MCF10A, 493 cells; MCF7, 550 cells; MDA-MB-231, 493 cells). Statistical significance is determined using two-way ANOVA followed by Sidak's multiple comparisons test. ***$p < 0.001$, ****$p < 0.0001$; ns, not significant. (B) Percentage of importin α1-positive MN in MCF10A, MCF7, and MDA-MB-231 cells with or without reversine treatment, quantified from indirect IF images stained with anti-importin α1 antibody. Data are shown as mean ± SD from three independent experiments, each including five microscopy fields (untreated: MCF10A, 95 MN; MCF7, 93 MN; MDA-MB-231, 102 MN; reversine: MCF10A, 100 MN; MCF7, 94 MN; MDA-MB-231, 90 MN). Statistical significance is determined using two-way ANOVA followed by Sidak's multiple comparison test. ns, not significant.

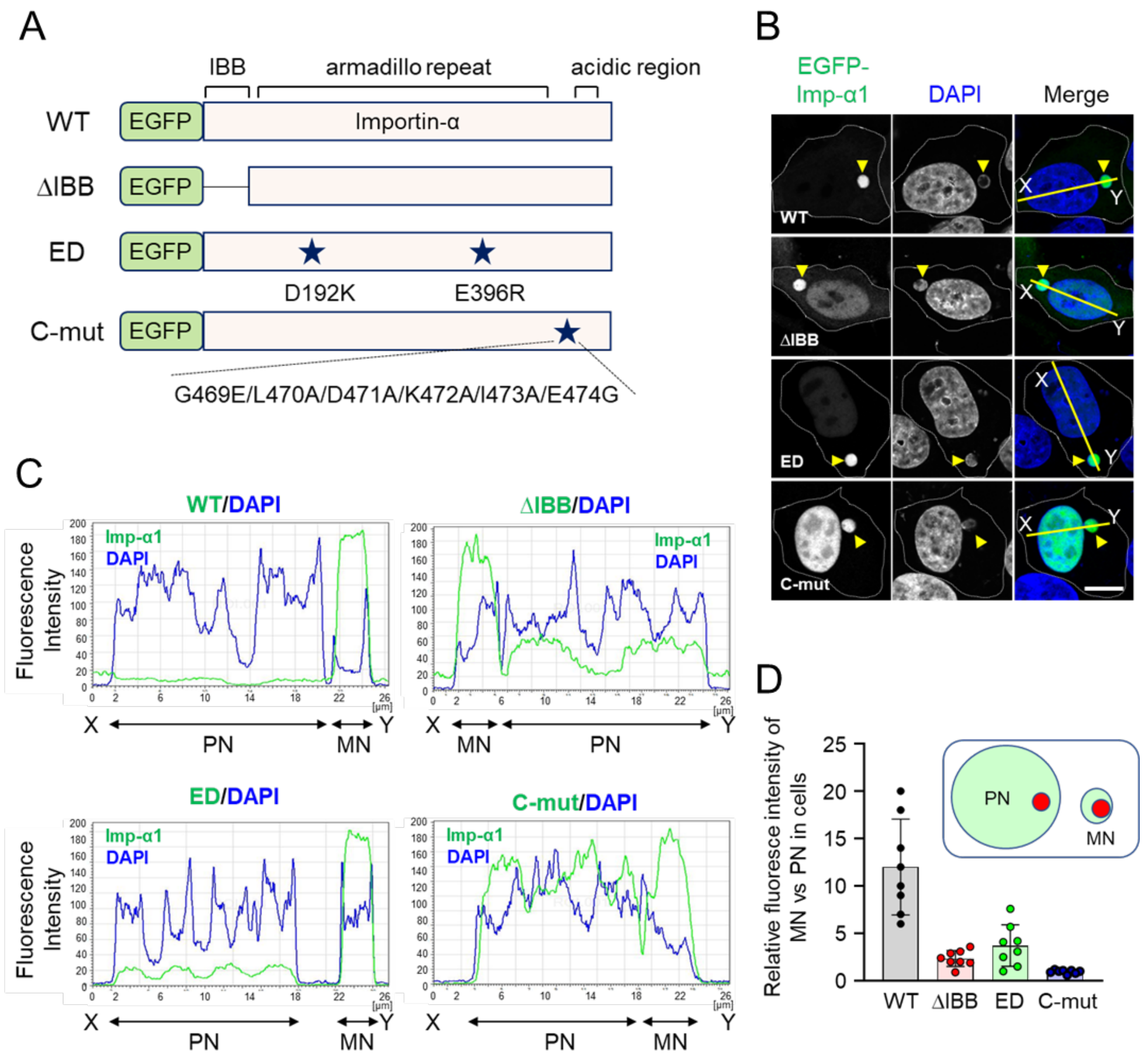


**Figure S4. Mutation analysis of importin α1 in MN localization**

(A) Schematic representation of importin α1 wild type (WT) and mutants: ΔIBB (deletion of the IBB domain), ED (NLS-binding–deficient, D192K/E396R), and C-mut (CAS-binding–defective, G469E/L470A/D471A/K472A/I473A/E474G). (B) Subcellular localization of EGFP–Imp-α1 WT or mutants in HeLa cells. Cells are transfected with EGFP constructs and treated with reversine to induce MN formation. White dotted lines outline the cellular boundaries. DNA is stained with DAPI. Yellow arrowheads indicate EGFP–Imp-α1-positive MN. The yellow X–Y lines described indicate the position used for fluorescence intensity analysis in panel C. (C) Fluorescence intensity profiles of EGFP-imp-α1 WT or mutants and DAPI along the X–Y line indicated in panel B. (D) Fluorescence intensity in a 2.0-μm diameter circle is measured in PN and MN in the same cell (see reference image, upper right; red circles indicate measurement area). Relative MN/PN values are plotted from eight independent cells.

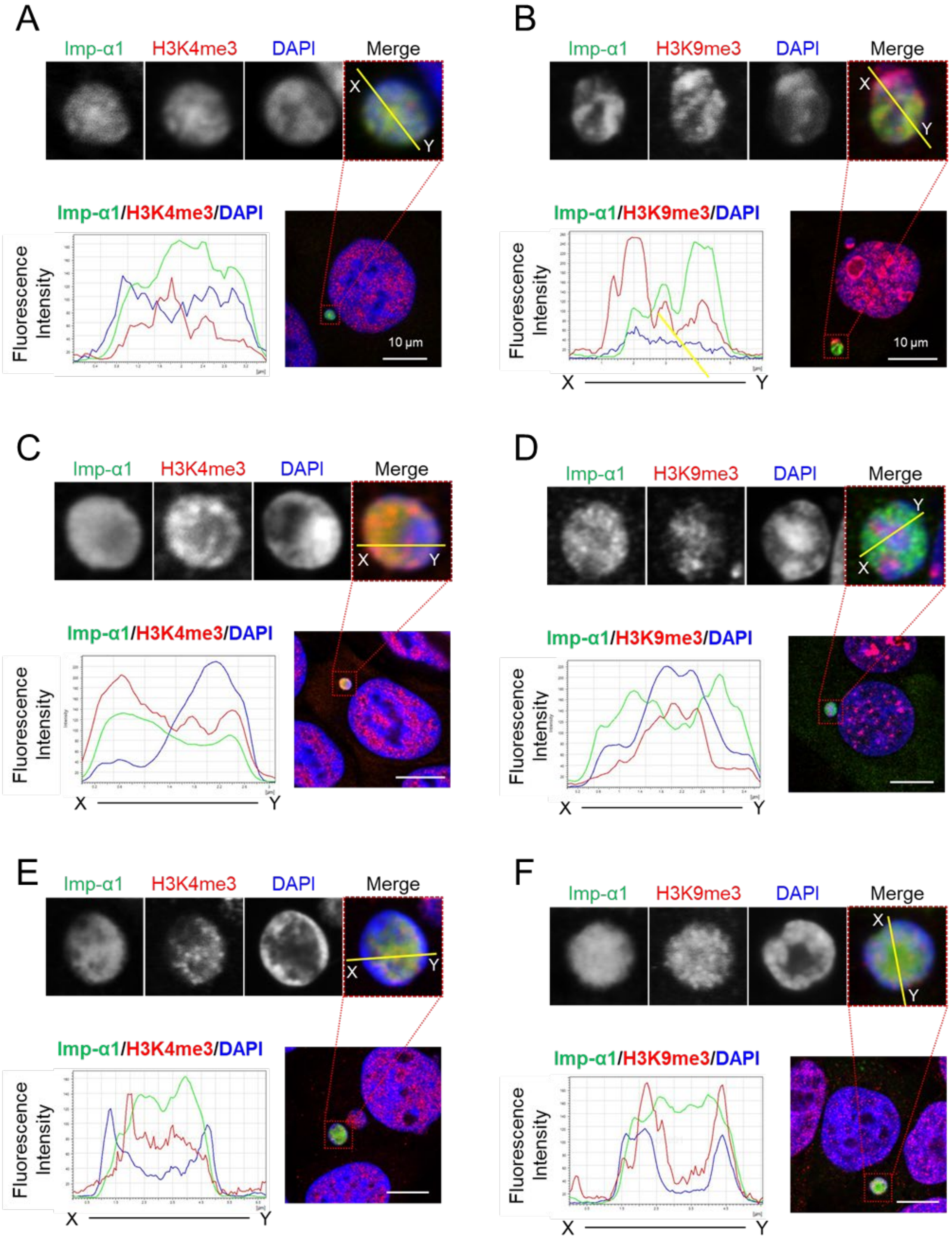


**Figure S5. Euchromatin localization of importin α1 in MN of human culture cell lines.**
(A–B) Indirect IF images of importin α1 with H3K4me3 (A) or H3K9me3 (B) in normally cultured MCF10A cells. The upper panels show magnified views of the areas indicated by the red dotted boxes in the lower panels. DNA is stained with DAPI. Fluorescent intensities of importin α1 (green), modified histones (red: H3K4me3 or H3K9me3), and DAPI (blue) in MN are plotted along the X–Y line and shown on the left. Scale bar: 10 µm. (C–D) Indirect IF images of importin α1 with H3K4me3 (C) or H3K9me3 (D) in normal cultured MCF7 cells.

The upper panels show magnified views of the areas indicated by red dotted boxes in the lower panels. DNA is stained with DAPI. Fluorescence intensities of importin α1 (green), modified histones (red: H3K4me3 or H3K9me3), and DAPI (blue) in MN are plotted along the X–Y line and shown on the left. Scale bar: 10 μm. (E–F) Indirect IF images of reversine-treated HeLa cells. The upper panels show magnified views of the areas indicated by the red dotted boxes in the lower panels. DNA is stained with DAPI. Fluorescent intensities of importin α1 (green), modified histones (red: H3K4me3 or H3K9me3), and DAPI (blue) in MN are plotted along the X–Y line and shown on the left. Scale bar: 10 μm.

**Table S1. Importin α1-binding candidate by RIME analysis**

| Gene Symbol | Gene Name | PANTHER Family/Subfamily (Panther number) | PANTHER Protein Class | Total Spectrum Count |
|---|---|---|---|---|
| ACTN1 | Alpha-actinin-1 | ALPHA-ACTININ-1 (PTHR11915:SF434) | actin or actin-binding cytoskeletal protein | 6 |
| ACTR2 | Actin-related protein 2 | ACTIN-RELATED PROTEIN 2 (PTHR11937:SF37) | actin and actin related protein | 6 |
| AP1B1 | AP-1 complex subunit beta-1 | AP-1 COMPLEX SUBUNIT BETA-1 (PTHR11134:SF3) | membrane traffic protein | 6 |
| ARF4 | ADP-ribosylation factor 4 | ADP-RIBOSYLATION FACTOR 4 (PTHR11711:SF110) | G-protein | 5 |
| ARHGDIA | Rho GDP-dissociation inhibitor 1 | RHO GDP-DISSOCIATION INHIBITOR 1 (PTHR10980:SF9) | G-protein modulator | 6 |
| BUB3 | Mitotic checkpoint protein BUB3 | MITOTIC CHECKPOINT PROTEIN BUB3 (PTHR10971:SF5) | RNA metabolism protein | 6 |
| BZW1 | Basic leucine zipper and W2 domain-containing protein 1 | BASIC LEUCINE ZIPPER AND W2 DOMAIN-CONTAINING PROTEIN 1 (PTHR14208:SF0) | basic leucine zipper transcription factor | 9 |
| CBX1 | Chromobox protein homolog 1 | CHROMOBOX PROTEIN HOMOLOG 1 (PTHR22812:SF159) | - | 6 |
| CCAR2/DBC1 | Cell cycle and apoptosis regulator protein 2 | CELL CYCLE AND APOPTOSIS REGULATOR PROTEIN 2 (PTHR14304:SF12) | chromatin/chromatin-binding, or -regulatory protein | 5 |
| CHD4 | Chromodomain-helicase-DNA-binding protein 4 | CHROMODOMAIN-HELICASE-DNA-BINDING PROTEIN 4 (PTHR45623:SF17) | chromatin/chromatin-binding, or -regulatory protein | 16 |
| DDX1 | ATP-dependent RNA helicase DDX1 | ATP-DEPENDENT RNA HELICASE DDX1 (PTHR24031:SF307) | RNA helicase | 8 |
| DDX23 | Probable ATP-dependent RNA helicase DDX23 | ATP-DEPENDENT RNA HELICASE DDX23-RELATED (PTHR47958:SF96) | RNA helicase | 6 |
| DHX15 | Pre-mRNA-splicing factor ATP-dependent RNA helicase DHX15 | PRE-MRNA-SPLICING FACTOR ATP-DEPENDENT RNA HELICASE DHX15 (PTHR18934:SF95) | RNA helicase | 8 |
| DNAJB1 | DnaJ homolog subfamily B member 1 | DNAJ HOMOLOG SUBFAMILY B MEMBER 1 (PTHR24078:SF568) | chaperone | 5 |
| EIF2S1 | Eukaryotic translation initiation factor 2 subunit 1 | EUKARYOTIC TRANSLATION INITIATION FACTOR 2 SUBUNIT 1 (PTHR10602:SF0) | translation initiation factor | 8 |
| EIF3A | Eukaryotic translation initiation factor 3 subunit A | EUKARYOTIC TRANSLATION INITIATION FACTOR 3 SUBUNIT A (PTHR14005:SF0) | translation initiation factor | 6 |
| EIF3I | Eukaryotic translation initiation factor 3 subunit I | EUKARYOTIC TRANSLATION INITIATION FACTOR 3 SUBUNIT I (PTHR19877:SF1) | translation initiation factor | 6 |
| FAM120A | Constitutive coactivator of PPAR-gamma-like protein 1 | CONSTITUTIVE COACTIVATOR OF PPAR-GAMMA-LIKE PROTEIN 1 (PTHR15976:SF14) | - | 14 |
| FEN1 | Flap endonuclease 1 | FLAP ENDONUCLEASE 1 (PTHR11081:SF58) | exodeoxyribonuclease | 8 |
| FKBP3 | Peptidyl-prolyl cis-trans isomerase FKBP3 | PEPTIDYL-PROLYL CIS-TRANS ISOMERASE FKBP3 (PTHR46493:SF1) | chaperone | 5 |
| FUS | RNA-binding protein FUS | RNA-BINDING PROTEIN FUS (PTHR23238:SF5) | RNA metabolism protein | 6 |
| G3BP1 | Ras GTPase-activating protein-binding protein 1 | RAS GTPASE-ACTIVATING PROTEIN-BINDING PROTEIN 1 (PTHR10693:SF21) | RNA metabolism protein | 5 |
| GTF2I | General transcription factor II-I | GENERAL TRANSCRIPTION FACTOR II-I (PTHR46304:SF2) | general transcription factor | 12 |
| HDAC1 | Histone deacetylase 1 | HISTONE DEACETYLASE 1 (PTHR48252:SF36) | histone modifying enzyme | 6 |
| HDAC2 | Histone deacetylase 2 | HISTONE DEACETYLASE 2 (PTHR48252:SF17) | histone modifying enzyme | 6 |
| HINT1 | Histidine triad nucleotide-binding protein 1 | HISTIDINE TRIAD NUCLEOTIDE-BINDING PROTEIN 1 (PTHR23089:SF45) | nucleotide phosphatase | 5 |
| HMGB2 | High mobility group protein B2 | HIGH MOBILITY GROUP PROTEIN B2 (PTHR48112:SF3) | chromatin/chromatin-binding, or -regulatory protein | 12 |
| HNRNPCL3 | Heterogeneous nuclear ribonucleoprotein C-like 3 | HETEROGENEOUS NUCLEAR RIBONUCLEOPROTEIN C-LIKE 1-RELATED (PTHR13968:SF30) | RNA metabolism protein | 13 |
| HNRNPUL2 | Heterogeneous nuclear ribonucleoprotein U-like protein 2 | HCG2044799-RELATED (PTHR12381:SF66) | - | 8 |
| ILF3 | Interleukin enhancer-binding factor 3 | INTERLEUKIN ENHANCER-BINDING FACTOR 3 (PTHR45762:SF4) | RNA metabolism protein | 11 |
| KPNA2 | Importin subunit alpha-1 | IMPORTIN SUBUNIT ALPHA-1 (PTHR23316:SF12) | transporter | 206 |
| LRRC47 | Leucine-rich repeat-containing protein 47 | LEUCINE-RICH REPEAT-CONTAINING PROTEIN 47 (PTHR10947:SF3) | aminoacyl-tRNA synthetase | 5 |
| LUC7L2 | Putative RNA-binding protein Luc7-like 2 | RNA-BINDING PROTEIN LUC7-LIKE 2-RELATED (PTHR12375:SF28) | - | 6 |
| MCM5 | DNA replication licensing factor MCM5 | DNA REPLICATION LICENSING FACTOR MCM5 (PTHR11630:SF42) | DNA metabolism protein | 8 |
| MCM6 | DNA replication licensing factor MCM6 | DNA REPLICATION LICENSING FACTOR MCM6 (PTHR11630:SF73) | DNA metabolism protein | 12 |
| MTA2 | Metastasis-associated protein MTA2 | METASTASIS-ASSOCIATED PROTEIN MTA2 (PTHR10865:SF4) | homeodomain transcription factor | 8 |
| MYO6 | Unconventional myosin-VI | UNCONVENTIONAL MYOSIN-VI (PTHR13140:SF745) | actin binding motor protein | 5 |
| NANS | Sialic acid synthase | SIALIC ACID SYNTHASE (PTHR42966:SF1) | acetyltransferase | 6 |

**Table S1 (continued)**

| | | | | |
|---|---|---|---|---|
| NAP1L1 | Nucleosome assembly protein 1-like 1 | NUCLEOSOME ASSEMBLY PROTEIN 1-LIKE 1 (PTHR11875:SF145) | chromatin/chromatin-binding, or -regulatory protein | 8 |
| NAP1L4 | Nucleosome assembly protein 1-like 4 | NUCLEOSOME ASSEMBLY PROTEIN 1-LIKE 4 (PTHR11875:SF75) | chromatin/chromatin-binding, or -regulatory protein | 8 |
| NCBP1 | Nuclear cap-binding protein subunit 1 | NUCLEAR CAP-BINDING PROTEIN SUBUNIT 1 (PTHR12412:SF2) | RNA splicing factor | 6 |
| NUDC | Nuclear migration protein nudC | NUCLEAR MIGRATION PROTEIN NUDC (PTHR12356:SF3) | microtubule or microtubule-binding cytoskeletal protein | 7 |
| NUP153 | Nuclear pore complex protein Nup153 | NUCLEAR PORE COMPLEX PROTEIN NUP153 (PTHR23193:SF23) | transporter | 10 |
| OPA1 | Dynamin-like 120 kDa protein, mitochondrial | DYNAMIN-LIKE 120 KDA PROTEIN, MITOCHONDRIAL (PTHR11566:SF67) | membrane traffic protein | 8 |
| PARP1 | Poly [ADP-ribose] polymerase 1 | POLY [ADP-RIBOSE] POLYMERASE 1 (PTHR10459:SF112) | DNA metabolism protein | 6 |
| PDAP1 | 28 kDa heat- and acid-stable phosphoprotein | 28 KDA HEAT- AND ACID-STABLE PHOSPHOPROTEIN (PTHR22055:SF5) | - | 7 |
| PDHB | Pyruvate dehydrogenase E1 component subunit beta, mitochondrial | PYRUVATE DEHYDROGENASE E1 COMPONENT SUBUNIT BETA, MITOCHONDRIAL (PTHR11624:SF96) | dehydrogenase | 5 |
| PMM2 | Phosphomannomutase 2 | PHOSPHOMANNOMUTASE 2 (PTHR10466:SF2) | mutase | 6 |
| PPM1G | Protein phosphatase 1G | PROTEIN PHOSPHATASE 1G (PTHR13832:SF321) | protein phosphatase | 6 |
| PPP1CA | Serine/threonine-protein phosphatase PP1-alpha catalytic subunit | SERINE/THREONINE-PROTEIN PHOSPHATASE PP1-ALPHA CATALYTIC SUBUNIT (PTHR11668:SF377) | protein phosphatase | 5 |
| PPP1R12A | Protein phosphatase 1 regulatory subunit 12A | PROTEIN PHOSPHATASE 1 REGULATORY SUBUNIT 12A (PTHR24179:SF20) | phosphatase modulator | 5 |
| PPP2R1A | Serine/threonine-protein phosphatase 2A 65 kDa regulatory subunit A alpha isoform | SERINE/THREONINE-PROTEIN PHOSPHATASE 2A 65 KDA REGULATORY SUBUNIT A ALPHA ISOFORM (PTHR10648:SF2) | phosphatase modulator | 6 |
| PRDX6 | Peroxiredoxin-6 | PEROXIREDOXIN-6 (PTHR43503:SF11) | peroxidase | 8 |
| PRPF3 | U4/U6 small nuclear ribonucleoprotein Prp3 | U4/U6 SMALL NUCLEAR RIBONUCLEOPROTEIN PRP3 (PTHR14212:SF0) | RNA splicing factor | 6 |
| PSMA5 | Proteasome subunit alpha type-5 | PROTEASOME SUBUNIT ALPHA TYPE-5 (PTHR11599:SF14) | protease | 5 |
| PSMD5 | 26S proteasome non-ATPase regulatory subunit 5 | 26S PROTEASOME NON-ATPASE REGULATORY SUBUNIT 5 (PTHR13554:SF10) | protease | 5 |
| PUF60 | Poly(U)-binding-splicing factor PUF60 | POLY(U)-BINDING-SPLICING FACTOR PUF60 (PTHR47330:SF1) | RNA splicing factor | 8 |
| PYGB | Glycogen phosphorylase, brain form | GLYCOGEN PHOSPHORYLASE, BRAIN FORM (PTHR11468:SF29) | glycosyltransferase | 8 |
| RACK1 | Receptor of activated protein C kinase 1 | RECEPTOR OF ACTIVATED PROTEIN C KINASE 1 (PTHR19868:SF0) | - | 6 |
| RANBP1 | Ran-specific GTPase-activating protein | RAN-SPECIFIC GTPASE-ACTIVATING PROTEIN (PTHR23138:SF135) | scaffold/adaptor protein | 6 |
| RANBP2 | E3 SUMO-protein ligase RanBP2 | E3 SUMO-PROTEIN LIGASE RANBP2-RELATED (PTHR23138:SF175) | scaffold/adaptor protein | 20 |
| RANGAP1 | Ran GTPase-activating protein 1 | RAN GTPASE-ACTIVATING PROTEIN 1 (PTHR24113:SF6) | GTPase-activating protein | 9 |
| RBM14 | RNA-binding protein 14 | RNA-BINDING PROTEIN 14 (PTHR23147:SF53) | RNA splicing factor | 10 |
| RGPD8 | RANBP2-like and GRIP domain-containing protein 8 | E3 SUMO-PROTEIN LIGASE RANBP2-RELATED (PTHR23138:SF175) | scaffold/adaptor protein | 6 |
| RPL13A | 60S ribosomal protein L13a | 60S RIBOSOMAL PROTEIN L13A (PTHR11545:SF30) | ribosomal protein | 11 |
| RPL18 | 60S ribosomal protein L18 | 60S RIBOSOMAL PROTEIN L18 (PTHR10934:SF6) | ribosomal protein | 10 |
| RPL19 | 60S ribosomal protein L19 | 60S RIBOSOMAL PROTEIN L19 (PTHR10722:SF34) | ribosomal protein | 5 |
| RPL21 | 60S ribosomal protein L21 | 60S RIBOSOMAL PROTEIN L21 (PTHR20981:SF8) | ribosomal protein | 6 |
| RPL26 | 60S ribosomal protein L26 | 60S RIBOSOMAL PROTEIN L26 (PTHR11143:SF11) | ribosomal protein | 6 |
| RPL27 | 60S ribosomal protein L27 | 60S RIBOSOMAL PROTEIN L27 (PTHR10497:SF11) | ribosomal protein | 7 |
| RPL27A | 60S ribosomal protein L27a | 60S RIBOSOMAL PROTEIN L27A (PTHR11721:SF27) | ribosomal protein | 8 |
| RPL35 | 60S ribosomal protein L35 | 60S RIBOSOMAL PROTEIN L35 (PTHR45722:SF9) | ribosomal protein | 8 |
| RPL35A | 60S ribosomal protein L35a | 60S RIBOSOMAL PROTEIN L35A (PTHR10902:SF30) | ribosomal protein | 8 |
| RPL37A | 60S ribosomal protein L37a | 60S RIBOSOMAL PROTEIN L37A (PTHR48160:SF1) | ribosomal protein | 6 |
| RPL5 | 60S ribosomal protein L5 | 60S RIBOSOMAL PROTEIN L5 (PTHR23410:SF24) | ribosomal protein | 8 |
| RPS15 | 40S ribosomal protein S15 | 40S RIBOSOMAL PROTEIN S15 (PTHR11880:SF28) | ribosomal protein | 6 |
| RPS23 | 40S ribosomal protein S23 | 40S RIBOSOMAL PROTEIN S23 (PTHR11652:SF44) | ribosomal protein | 5 |

**Table S1 (continued)**

| | | | | |
|---|---|---|---|---|
| RPS24 | 40S ribosomal protein S24 | 40S RIBOSOMAL PROTEIN S24 (PTHR10496:SF7) | ribosomal protein | 6 |
| RPS5 | 40S ribosomal protein S5 | 40S RIBOSOMAL PROTEIN S5 (PTHR11205:SF34) | ribosomal protein | 6 |
| RTCB | tRNA-splicing ligase RtcB homolog | TRNA-SPLICING LIGASE RTCB HOMOLOG (PTHR11118:SF1) | - | 8 |
| S100A11 | Protein S100-A11 | PROTEIN S100-A11 (PTHR11639:SF60) | calmodulin-related | 9 |
| SARS | Serine--tRNA ligase, cytoplasmic | SERINE--TRNA LIGASE, CYTOPLASMIC (PTHR11778:SF7) | aminoacyl-tRNA synthetase | 6 |
| SART3 | Squamous cell carcinoma antigen recognized by T-cells 3 | SQUAMOUS CELL CARCINOMA ANTIGEN RECOGNIZED BY T-CELLS 3 (PTHR15481:SF5) | RNA splicing factor | 12 |
| SF3A2 | Splicing factor 3A subunit 2 | SPLICING FACTOR 3A SUBUNIT 2 (PTHR23205:SF0) | RNA splicing factor | 5 |
| SH3GLB2 | Endophilin-B2 | DREBRIN-LIKE PROTEIN-RELATED (PTHR14167:SF68) | scaffold/adaptor protein | 6 |
| SMARCA5 | SWI/SNF-related matrix-associated actin-dependent regulator of chromatin subfamily A member 5 | SWI/SNF-RELATED MATRIX-ASSOCIATED ACTIN-DEPENDENT REGULATOR OF CHROMATIN SUBFAMILY A MEMBER 5 (PTHR10799:SF997) | DNA helicase | 6 |
| SNRNP70 | U1 small nuclear ribonucleoprotein 70 kDa | U1 SMALL NUCLEAR RIBONUCLEOPROTEIN 70 KDA (PTHR13952:SF5) | RNA splicing factor | 6 |
| SRRM2 | Serine/arginine repetitive matrix protein 2 | SERINE/ARGININE REPETITIVE MATRIX PROTEIN 2 (PTHR34755:SF3) | - | 14 |
| STARD10 | PCTP-like protein | PCTP-LIKE PROTEIN (PTHR19308:SF7) | - | 6 |
| SUPT16H | FACT complex subunit SPT16 | FACT COMPLEX SUBUNIT SPT16 (PTHR13980:SF17) | chromatin/chromatin-binding, or -regulatory protein | 8 |
| TBCB | Tubulin-folding cofactor B | TUBULIN-FOLDING COFACTOR B (PTHR18916:SF85) | chaperone | 6 |
| THRAP3 | Thyroid hormone receptor-associated protein 3 | THYROID HORMONE RECEPTOR-ASSOCIATED PROTEIN 3 (PTHR15268:SF16) | DNA metabolism protein | 6 |
| TOP2B | DNA topoisomerase 2-beta | DNA TOPOISOMERASE 2-BETA (PTHR10169:SF36) | DNA metabolism protein | 16 |
| TPR | Nucleoprotein TPR | NUCLEOPROTEIN TPR (PTHR18898:SF2) | primary active transporter | 6 |
| TRA2B | Transformer-2 protein homolog beta | TRANSFORMER-2 PROTEIN HOMOLOG BETA (PTHR48034:SF1) | RNA metabolism protein | 6 |
| TRIM33 | E3 ubiquitin-protein ligase TRIM33 | E3 UBIQUITIN-PROTEIN LIGASE TRIM33 (PTHR45915:SF3) | chromatin/chromatin-binding, or -regulatory protein | 8 |
| UBA1 | Ubiquitin-like modifier-activating enzyme 1 | UBIQUITIN-LIKE MODIFIER-ACTIVATING ENZYME 1 (PTHR10953:SF155) | ubiquitin-protein ligase | 8 |
| UNC45A | Protein unc-45 homolog A | PROTEIN UNC-45 HOMOLOG A (PTHR45994:SF3) | - | 8 |
| UPF1 | Regulator of nonsense transcripts 1 | REGULATOR OF NONSENSE TRANSCRIPTS 1 (PTHR10887:SF364) | RNA helicase | 10 |
| USP5 | Ubiquitin carboxyl-terminal hydrolase 5 | UBIQUITIN CARBOXYL-TERMINAL HYDROLASE 5 (PTHR24006:SF655) | cysteine protease | 12 |
| USP7 | Ubiquitin carboxyl-terminal hydrolase 7 | UBIQUITIN CARBOXYL-TERMINAL HYDROLASE 7 (PTHR24006:SF644) | cysteine protease | 6 |
| ZNF185 | Zinc finger protein 185 | ZINC FINGER PROTEIN 185 (PTHR15468:SF2) | - | 6 |

**Table S2. MN localization with Importin α1**

| Mapped IDs | Gene Name, Gene Symbol | % vs all MN | % overlap with Imp-α1 |
|---|---|---|---|
| CCAR2/DBC1 | Cell cycle and apoptosis regulator protein 2 | 63% | 95% |
| TOP2B | DNA topoisomerase 2-beta | 79% | 93% |
| SUPT16H | FACT complex subunit SPT16 | 68% | 92% |
| PARP1 | Poly [ADP-ribose] polymerase 1 | 92% | 91% |
| GTF2I | General transcription factor II-I | 100% | 75% |
| HDAC1 | Histone deacetylase 1 | 100% | 67% |
| HDAC2 | Histone deacetylase 2 | 100% | 63% |
| DNAJB1 | DnaJ homolog subfamily B member 1 | 66% | 52% |

**Table S3. Primary antibody details**

| Name in paper | | Source | Catalogue # | Species |
|---|---|---|---|---|
| Importin α5/KPNA1 | | Santa Cruz | sc-6918 | goat |
| | | Abnova | 2A4-1B5 | mouse |
| Importin α1/KPNA2 | Ab1 | BD Transduction. | 610486 | mouse |
| | Ab2 | Abcam | ab84440 | rabbit |
| | Ab3 | Abcam | ab6036 | goat |
| Importin α4/KPNA3 | | Imgenex | IMG3568 | goat |
| Importin α3/KPNA4 | | Abcam | ab6039 | goat |
| Importin α6/KPNA5 | | Supplemental materials and methods | 1E3-2D6 | mouse |
| Importin α7/KPNA6 | | Cell Engineering Co. | 3F8 | rat |
| Importin β1/KPNB1 | | Santa Cruz | sc-1919 | goat |
| | | Abcam | ab2811 | mouse |
| CAS/CSE1L | | Abcam | ab96755 | rabbit |
| Lamin A/C | | Santa Cruz | sc-6215 | goat |
| Lamin B1 | | MBL | PM064 | rabbit |
| H3K4me3 | | MBL | MABI0304 | mouse |
| | | Supplemental materials and methods | CMA304 | mouse |
| H3K9me3 | | MBL | MABI0308 | mouse |
| | | Supplemental materials and methods | CMA318 | mouse |
| RAD51 | | Abcam | ab133534 | rabbit |
| MAb414 | | Biolegend | MMS-120P | mouse |
| ACTIN | | Santa Cruz | sc-1615 | goat |
| Flag (M2) | | Sigma-aldrich | F1804 | mouse |
| CCAR2/DBC1 | | CST | 5857 | mouse |
| TOP2B | | SIGMA | HPA024120 | rabbit |
| SUPT16H | | SIGMA | HPA049787 | rabbit |
| PARP1 | | Proteintech | 66520-1-Ig | mouse |
| GTF2I | | SIGMA | HPA026638 | rabbit |
| HDAC1 | | CST | 5356 | mouse |
| HDAC2 | | CST | 5113 | mouse |
| DNAJB1 | | Proteintech | 13174-1-AP | rabbit |
| RPA2/RPA32 | | CST | 2208 | rat |
| cGAS | | CST | 79978 | rabbit |
